# Memory retrieval explains dynamic effects of expectations on perceptual decisions

**DOI:** 10.64898/2026.09.22.753289

**Authors:** Ari Khoudary, Megan A. K. Peters, Aaron M. Bornstein

**Author notes:** Corresponding author: Ari Khoudary. These authors contributed equally (Megan A. K. Peters, Aaron M. Bornstein). Conflicts of Interest: M.A.K.P. is a consultant for the for-profit entity Conscium, Inc., which seeks to pioneer safe, efficient artificial intelligence and which played no role in this project’s conceptualization, analyses, interpretation, or writing. The authors declare no conflicts of interest.

## Abstract

Expectations—prior knowledge of environmental statistics—support adaptive behavior by reducing uncertainty during inference. However, the questions of *where* expectations come from and how that *ought* to impact their integration has remained largely unexplored. Here, we present a novel model formalizing the role of *memory retrieval* in the process of integrating expectations with uncertain sensory information. To do this, we first articulate how *uncertainty about expectations*—omitted from existing normative models—ought to induce memory retrieval processes. We then model expectations as a dynamic source of internal evidence that observers sample from *in parallel* to vision, and that are weighed according to a time-evolving estimate of their reliability *relative* to sensory evidence. Finally, we show that this single, principled model can reproduce several previously disparate observations of *time-varying* effects of expectations in neural and behavioral data, each of which was previously explained using different neural and computational mechanisms. These findings suggest that our model identifies a basic mechanism governing the integration of expectations with sensory evidence on assumptions that more closely approximate the dynamic, heterogeneous, and uncertain nature of real-world decision settings.

## 1 Introduction

In order to stay alive, humans and other animals must make quick decisions about how to act on the basis of noisy, ambiguous, and limited sensory information. Expectations—prior knowledge about statistical regularities—are a powerful source of information that decision-makers use to reduce uncertainty and make more efficient decisions (e.g., Seriès and Seitz, 2013; Summerfield and de Lange, 2014; de Lange et al., 2018). The decades of neural and behavioral evidence supporting this claim rest upon normative formal models that describe the optimal procedure for combining expectations with uncertain sensory evidence (Wald and Wolfowitz, 1948; Edwards, 1965; Bogacz et al., 2006; Simen et al., 2009; Moran, 2015; Malhotra et al., 2018). Critically, however, the assumptions of these normative models have systematically obscured investigation into how *uncertainty* about expectations modulates their integration into perceptual decisions. This conceptual gap is especially notable because of the ubiquity of uncertain expectations in decisions made outside of laboratory settings, where observers do not have information about *which* expectation to use and/or *what* its true predictive probability is.

To address this gap, we develop a computational model describing a principled solution to the problem of combining uncertain expectations with sensory evidence that is itself uncertain. Using the formalism of sequential sampling, we posit that uncertainty about expectations induces dynamic retrieval processes that continuously sample information from memory to reduce uncertainty in the expectation’s prediction. Critically, we posit that this retrieval process happens *in parallel* to sensory evidence sampling, such that two dynamic evidence signals enter into the decision process. Drawing on cue combination and multisensory integration, we then model the key quantity driving choice—the time-evolving decision variable—as a dynamic, reliability-weighted sum of samples from the parallel memory and sensory evidence timeseries. This creates an adaptive decision process that is sensitive to moment-by-moment fluctuations in the informativeness of sensory evidence, dynamically increasing reliance on expectations only when sensory evidence is too noisy to reliably guide behavior.

Then, to support the theoretical and empirical validity of our model, we present a series of simulation results, capturing data observed in different species, task designs, and neuroimaging modalities. Our simulation findings both demonstrate that dynamic reliability-weighted integration offers a unifying explanation for these previously-disparate observations, and provide empirical evidence for one of the key latent quantities in our model: a dynamic signal of how much weight to place on expectations *within* the timecourse of a single choice. Taken together, this work challenges traditional static models of expectations in perceptual decision-making, suggesting instead that the dynamics of memory retrieval—functioning to reduce uncertainty in the expectation—generate time-varying effects on perceptual decisions.

## 2 Static effects of expectations on perceptual decisions: formal models and empirical data

To begin, we review the origins and development of static models of expectations in perceptual decision-making. We first describe the sequential sampling framework within which all models in this article are specified, and then introduce the diffusion decision or drift-diffusion model (DDM). We focus on this model because it is the foundation upon which static models of expectations were initially developed. To motivate the form of our own dynamic model, we discuss how static models have been implemented, what they assume, and the neural and behavioral evidence supporting differently-formalized static (i.e., time-constant) effects of expectations.

### 2.1 Sequential sampling and the diffusion decision model (DDM) of two-alternative choice

The most common framework for studying perceptual decisions based on dynamically-evolving sensory evidence is *sequential sampling*, often termed evidence accumulation (Gold and Shadlen, 2007; Forstmann et al., 2016; Ratcliff et al., 2016). This conceptual framework for studying the psychology and neuroscience of (mainly) two-alternative decisions is based on the mathematical framework of sequential analysis in statistics (Barnard, 1946; Wald, 1945; Gold and Shadlen, 2002). Both frameworks posit that decisions are made by continuously sampling information from an evidence source, extracting and integrating decision-relevant information over time, and committing to one of the options once the accumulated evidence surpasses a threshold value. The formal framework of sequential analysis yields normative solutions to the accumulation process, and thus can be used to build models that optimize the tradeoff between decision accuracy and deliberation time (the *speed-accuracy tradeoff*).

Importantly, not all sequential sampling models are normative. Some can be proven to non-optimally trade off speed and accuracy (e.g., race models; Vickers, 1970; Bogacz et al., 2006), whereas the normativity of others cannot be evaluated at all because of their mathematical form (e.g., the extended diffusion decision model; Ratcliff and Tuerlinckx, 2002; Bogacz et al., 2006). In an influential paper, Bogacz et al. (2006) demonstrated that the earliest form of the *diffusion decision model* (or drift-diffusion model; DDM) optimally trades off speed with accuracy under different formalizations of that trade-off. Additionally, they showed that five other prominent models become equivalent to the DDM when their parameter settings optimize those same criteria (Bogacz et al., 2006), providing strong evidence for the normativity of the process specified by the DDM. Because early starting point models were designed to retain the optimality of the DDM while still incorporating a bias toward one option (Edwards, 1965; Link, 1975; Bogacz et al., 2006), we turn next to describing the DDM.

The first application of a sequential sampling modeling to the study of human decision processes is attributed to Stone (1960), who posited that the dynamics of two-alternative deliberation could be captured by a Gaussian random walk:

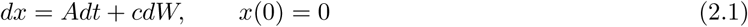

where *x*(0) represents the starting point of the decision variable, *dx* represents a change in the decision variable *x* over a unit of time *dt*, *A* represents the internal signal strength (corresponding to the drift rate of the process), and *cdW* represents noise/diffusion in the accumulation process which follows a normal distribution with mean 0 and variance *c*^2^*dt* (Bogacz et al., 2006). The process ends once the decision variable surpasses a threshold value *z*, at which point the decision maker commits to the choice corresponding to the threshold value reached (+*z* or −*z*). Throughout, we will call this model the “original” DDM to distinguish it from the “extended” DDM which is a probabilistic extension designed to maximize the model’s ability to fit behavior across a wide range of experimental conditions (Ratcliff and Rouder, 1998; Ratcliff and Tuerlinckx, 2002; Ratcliff et al., 2016).

The optimality of the original DDM comes from its equivalence with the sequential probability ratio test (SPRT; Bogacz et al., 2006; Moran, 2015). The SPRT gives the optimal procedure for determining when enough evidence has been gathered to terminate a two-alternative, discrete sampling process (Wald, 1945; Barnard, 1946; Wald and Wolfowitz, 1948). Optimality on the SPRT (hereafter “SPRT-optimality”) is defined as minimizing the average amount of time needed to make a decision at a fixed level of accuracy, or maximizing the proportion of correct responses for a fixed decision time (Wald and Wolfowitz, 1948; Bogacz et al., 2006). We discuss this form of optimality and its relationship to other forms at length in Khoudary et al. (2025b); a comprehensive overview is also provided by Bogacz et al. (2006).

Finally, it is worth mentioning that, on both the original and extended DDM, the drift rate *A* and prior probability Π terms are assumed to be functions of the decision environment. The drift rate term *A* has been robustly shown to be proportional to the signal-to-noise ratio in sensory stimuli (often termed the *coherence* of evidence; Palmer et al., 2005a; Gold and Shadlen, 2007; Ratcliff et al., 2016). The prior probability term Π likewise has been shown to scale with the frequency at which each choice outcome is presented (often termed the *bias* in the environment; Palmer et al., 2005b; Simen et al., 2009; Moran, 2015). The diffusion noise term *c*^2^ is most commonly treated as a fixed parameter during model fitting and is not commonly linked to properties of the decision environment (though see Zylberberg et al., 2016). The threshold term *z* is *optimally* defined as a function of the decision environment, but *psychologically* corresponds to individual differences in risk tolerance (Bogacz et al., 2006; Ratcliff et al., 2016), and thus is commonly allowed to vary across individuals during model fitting. In what follows, we focus on normative considerations for the drift rate *A* and prior probability Π terms. Because SPRT assumes a time-constant decision boundary, we assume the same for all discussions that follow.

### 2.2 Starting point models of expectations

The most prominent approach to incorporating expectations into sequential sampling models—and the DDM specifically—is adjusting the starting point *x*(0) of the decision process in proportion to the magnitude of the expectation (Edwards, 1965; Bogacz et al., 2006; Simen et al., 2009; Gold and Shadlen, 2007, Ratcliff et al., 2016). This section reviews the normative principles motivating this convention, the assumptions upon which its optimality rests, and empirical evidence supporting starting point formalizations of expectations.

#### 2.2.1 Normative grounding for starting point models

Edwards (1965) proposed one of the first extensions to the original DDM: adjusting the starting point of the decision process *x*(0) to capture any existing biases that an agent might have toward deciding in favor of one of the two outcomes. Let Π denote the probability with which the choice option corresponding to +*z* is correct on each trial. An *environment* is considered biased if Π ̸= 0.5; i.e., if one of the two options is more frequently correct or rewarded relative to the other. Conversely, an *observer* is inferred to be biased toward one of the two options if Π = 0.5 but their choices are best described by a model where *x*(0) ̸= 0. Edwards (1965) showed that the optimality of the original DDM can be retained if, in a biased environment, observers adjust their starting point *x*(0) according to:

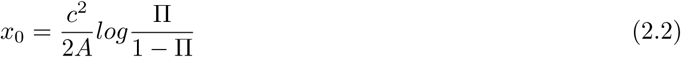

where *A* is the drift rate and *c* is the variance of a Gaussian diffusion process (Bogacz et al., 2006). A highly similar model was proposed by Link (1975), who instead posited that the log-odds ratio be scaled by a factor of 4. These differences stem from different formalizations of the speed-accuracy tradeoff, which we discuss in the next section. The *similarities* in the models, however, reveal two general principles about how expectations are optimally integrated into perceptual decisions. First, the starting point should be offset in proportion to the log-odds ratio of choice outcomes before observing any evidence. Second, this offset should be scaled by the *ratio* of noise (*c*^2^) to signal (*A*) in the decision environment (Bogacz et al., 2006). Critically, this scaling factor leads to exponentially greater weighting of expectations in decisions as the signal-to-noise ratio of evidence approaches zero (Figure **??**). In other words, expectations ought to be weighted more strongly as the uncertainty of sensory evidence increases.

#### 2.2.2 Assumptions and limitations of the normative starting point model

The small difference in the forms of Edwards’ (1965) and Link’s (1975) models stem from different choices about how to formalize the speed-accuracy tradeoff as an optimality criterion. Edwards 1965 used the standard definition of SPRT-optimality (Section 2.1), whereas Link’s (1975) model minimizes error rate for a fixed decision threshold. Although these different definitions of optimality led to nearly identical solutions in this case, different definitions of optimality lead to qualitatively different normative models (and, by extension, distinct theories of optimal decision processes). A core argument of this article is that the normativity of the starting point model—along with its empirical utility—has overly simplified formal theories of expectation-setting and integration. Specifically, we propose that formal models and experimental settings alike have largely obscured the role that *uncertainty* about expectations has in shaping how they are integrated with sensory evidence that is itself uncertain (Figure 1). Incorporating uncertainty about expectations is important not only for capturing real-world scenarios within which observers make decisions, but also for interpreting the dynamics of information transfer across brain regions in more complex decision-making settings (e.g., Hachisuka, Shor, Liu et al., 2026).

**Figure 1:**
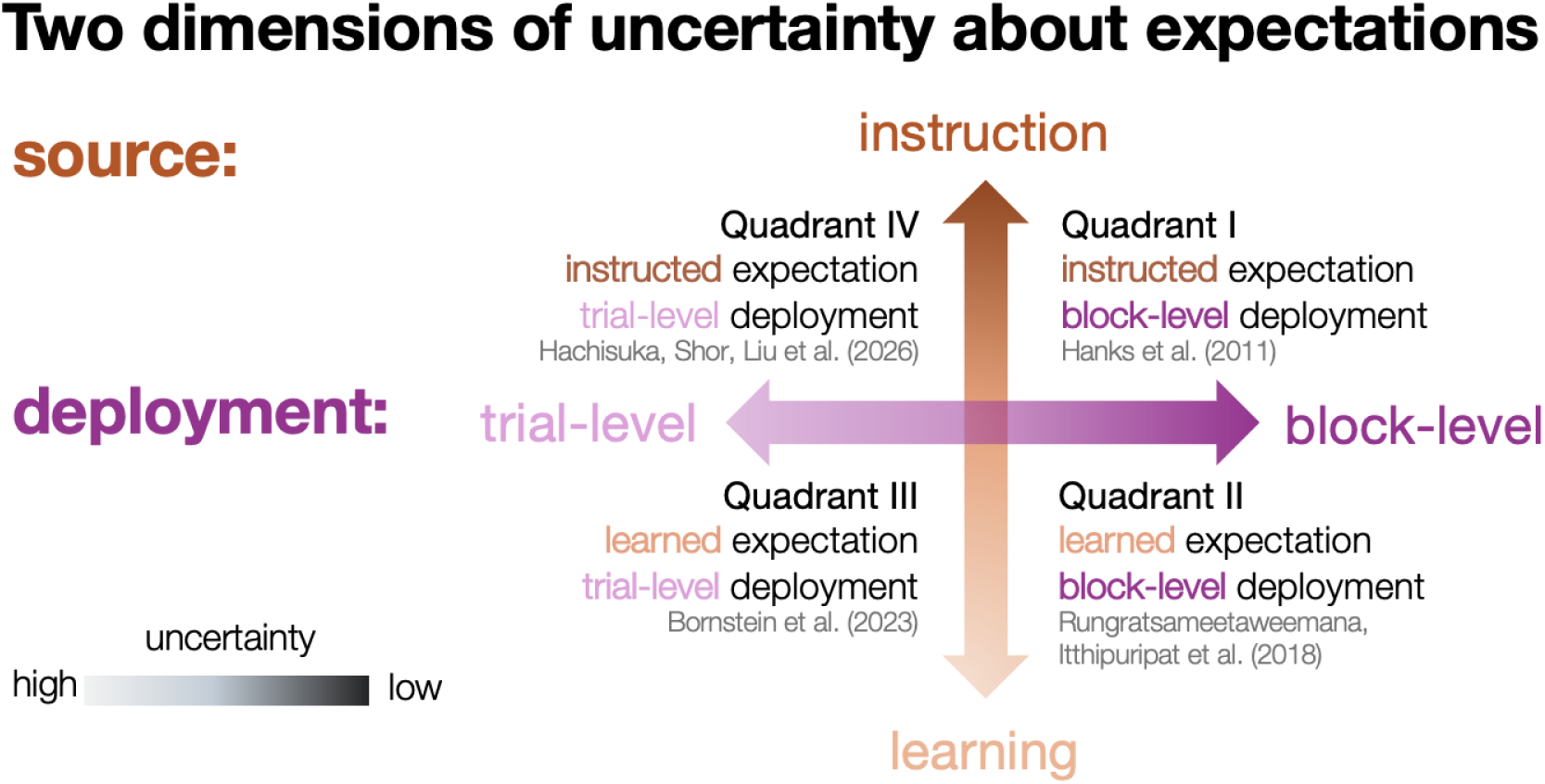
Two dimensions of uncertainty that observers can have about expectations when making perceptual decisions. Source-dependent uncertainty (orange) captures uncertainty about *what* an expectation predicts, whereas deployment-dependent uncertainty (pink) captures uncertainty about *which* expectation to use for each decision. The relative degree of uncertainty within a dimension is represented by the opacity of its respective color, with highly uncertain conditions in lighter shades and less certain conditions in darker shades. Gray-text citations indicate existing findings of dynamic effects observed in each quadrant; we show that our model can capture each of these findings in Section 5.

To demonstrate these points, we begin by discussing the two assumptions that models must meet in order to be considered SPRT-optimal, and how these assumptions are violated in settings of considerable scientific interest. Assumption 1 describes conditions the decision environment must meet, whereas Assumption 2 describes conditions that the observer’s knowledge must meet in order to execute the SPRT-optimal procedure. In this way, SPRT-optimality can be thought to be comprised of conditions on both external (Assumption 1) and internal (Assumption 2) factors of the decision process.

##### 2.2.2.1 Assumption 1: homogeneous decision environment

The first assumption that any SPRT-optimal model must make is that properties of the decision environment are *fixed*, *constant*, or *homogeneous*. This means that the drift rate *A*, diffusion magnitude *c*^2^, and prior probability Π *should* take on one scalar value that is constant across all decisions made in that context. On this assumption, any trial-level variability in the accumulation process is attributed solely to the noise term *c*^2^. When experimental conditions are deliberately constructed such that this first assumption of SPRT-optimality is met, behavior in humans and non-human animals appears largely SPRT-optimal (Link, 1975; Bogacz et al., 2006; Simen et al., 2009; Brunton et al., 2013).

However, the utility of SPRT-optimality for reasoning normatively about human behavior is limited for at least two reasons. First, real-world choice environments hardly—if ever—are homogeneous on all the dimensions that SPRT-optimality requires. Second, experimental tasks are also rarely homogeneous in an SPRT-compatible way. Unless an experiment explicitly sets out to test a prediction of an SPRT-optimal model, it is likely to manipulate the signal strength or difficulty of a decision at least pseudorandomly across trials. An important implication, then, is that arguments for starting point models that appeal to optimality are only valid for behavioral data collected in homogeneous decision environments. Once some trial-level variability in the signal strength *A*, evidence volatility *c*^2^, or prior probability Π is introduced, the starting point model is no longer the optimal procedure for integrating expectations about choice outcomes into the decision process (Moran, 2015).

##### 2.2.2.2 Assumption 2: full access to properties of the environment

The second assumption required for SPRT-optimality is that decision-makers have access to the fixed, true values of the environmental properties corresponding to the parameters *A*, *c*^2^, and Π: sensory signal strength, sensory signal volatility, and the prior probability of either choice option. This assumption follows directly from the procedure on which SPRT optimizes the speed-accuracy tradeoff. An observer who has access to the true values of *A*, *c*^2^, and Π, along with their desired proportion of correct responses, can, in theory, compute the precise threshold value *z* that leads to SPRT-optimal behavior (minimizing the average amount of time needed to respond at a desired level of accuracy).

A core limitation on the utility of this assumption is that humans and other animals rarely—if ever—have direct access to this information when making perceptual decisions in the real world. Instead, multiple learning and memory systems work together to store and flexibly utilize information about statistical regularities, both about the decision-maker and their environment, acquired across different timescales of experience (Seriès and Seitz, 2013; de Lange et al., 2018; Wang et al., 2021; Bakkour et al., 2019; Yoo and Bornstein, 2024; Gläscher et al., 2010; Bornstein and Daw, 2013, 2012; Noh et al., 2023). A more realistic starting premise, then, is that decision-makers *learn* about the values of *A*, *c*^2^, and Π through experience and — critically—treat these sparsely-informative expectations as appropriately uncertain. Accordingly, observers must rely on their learning processes to optimize behavior, while *also* incorporating their uncertainty about the learned values into the optimization process (e.g., Hanks et al., 2011; Khodadadi et al., 2014; Findling et al., 2025). We expand upon these points in Sections 2.3.2.2 and 3.2.1.

#### 2.2.3 Empirical support for starting point models

A number of empirical findings support the idea that expectations bias perceptual decisions by modifying the starting point of the evidence accumulation process (Carpenter and Williams, 1995; Ratcliff and McKoon, 2008; Simen et al., 2009; van Ravenzwaaij et al., 2012; Mulder et al., 2012; de Lange et al., 2013; Bornstein et al., 2023). Early work initially identified starting point biases via fits to human behavior (Carpenter and Williams, 1995, Ratcliff and McKoon, 2008, Simen et al., 2009, van Ravenzwaaij et al., 2012), with later work linking starting point effects to neural activity in humans (de Lange et al., 2013, Mulder et al., 2012, Bornstein et al., 2023) and non-human primates (Rao et al., 2012). This section briefly reviews these findings, which demonstrate that (1) humans do appear to optimally integrate expectations with sensory evidence and (2) expectations exert effects on the starting point even when the conditions for SPRT-optimality are not met.

One of the few experiments that explicitly created an environment that meets the assumptions for SPRT-optimality was conducted by Simen et al. (2009). The authors fixed signal strength *A* across trials, and manipulated the prior probability of choice outcomes Π across blocks of trials. In support of optimal behavior, the authors found selective effects of prior probability Π on the starting point of the decision process: starting point values were selectively greater in the biased blocks than in the neutral blocks (Simen et al., 2009). The authors additionally manipulated the amount of trials—and the time elapsed between them—across blocks, such that some blocks contained several trials presented quickly in succession, whereas other blocks contained fewer trials with greater amounts of time elapsed between them. Combining this manipulation with that of the prior probability makes a somewhat surprising prediction: as the amount of elapsed time between trials becomes sufficiently low, and the prior probability Π becomes sufficiently high, observers should opt for a *non-integrative* decision procedure: responding immediately with the choice that has a higher Π (Simen et al., 2009).

This prediction follows directly from (1) the optimality of lowering thresholds for faster-paced choice environments with more opportunities to make a decision and (2) the optimality of biasing the starting point toward the more likely option in proportion to Π.^1^ RT distributions across conditions demonstrated that observers indeed increasingly adopted non-integrative strategies: at the highest levels of Π and task-pacing, most responses were made immediately at the onset of sensory evidence (Simen et al., 2009). Conversely, as the pacing of the task slowed down—and observers had fewer opportunities to make correct responses— the proportion of non-integrative responses *decreased*, an effect that is particularly pronounced in the most biased environments (Π = 0.9; Simen et al., 2009). Taken together, these findings suggest that humans can optimally modify their decision processes to account for temporal and statistical properties of different choice environments.

A study by Mulder et al. (2012) used functional neuroimaging to further investigate the neural correlates of starting point effects. The authors contrasted two types of expectations: one consisting of information about the likely outcome (i.e., prior probability) and another consisting of information about the outcome more likely to be rewarded (i.e., potential payoff). To do this, they incorporated a cue at the start of each trial of a motion discrimination task. The cues consisted of a left- or right-pointing arrow and either a “%” (indicating prior probability) or “€” (indicating potential payout). Although this trial-level manipulation of expectations makes the SPRT no longer optimal, Mulder et al. (2012) found that both types of cues induced effects on the starting point. Further, they found that starting point offsets returned by the model parametrically modulated BOLD activity in frontoparietal regions and anterior cingulate gyrus, suggesting these areas may encode information about prior probability during decision-making (Mulder et al., 2012).

Finally, a study by Rao et al. (2012) revealed neural evidence of a starting point effect in rhesus macaques performing a motion discrimination task. The authors obtained electrophysiological measurements from neurons in measurements from neurons in cortical areas previously shown to dissociably encode sensory and decisional information (middle temporal (MT) and lateral intraparietal cortex (LIP), respectively). Rao et al., 2012 found that trial-varying cues induce directional activity in LIP, such that firing rates are increased for neurons selective to the cued motion direction and decreased for neurons selective to the alternative. The authors did not find an effect of expectations on firing rates in MT, which led them to conclude that expectations modulate decisions by selectively biasing the starting point of evidence.

Taken together, the work reviewed in this section lends empirical support for a starting point effect of expectations. Model fitting to human behavior reveals that starting point biases describe effects of expectations well, further suggesting that human perceptual decision-making is largely optimal. Neuroimaging work has linked these starting point effects to neural activity in frontoparietal regions, suggesting that expectations enter into the decision process as an additional source of evidence that’s integrated with sensory information to optimize the decision process. However, a growing body of work suggests that expectations might *also* exert effects on the drift rate of the decision process (Hanks et al., 2011; Moran, 2015; Dunovan and Wheeler, 2018; Diaz et al., 2024; Kelly et al., 2021; Bornstein et al., 2023). We discuss these findings and how they relate to optimal decision models in the next section.

### 2.3 Drift rate models of expectations

Whereas starting point models posit that expectations simply shift the initial value of the decision process *x*(0), drift rate models posit that the time-evolving evidence itself is accumulated more efficiently for outcomes that have a higher prior probability Π. Conceptually, drift rate models posit that expectations *amplify* the signal of expectation-congruent evidence and *attenuate* the signal of expectation-incongruent evidence. This type of effect on decisions is qualitatively similar to models that posit a sharpening of sensory representations as a function of expectations (e.g., Kok et al., 2012; Afacan-Seref et al., 2018), but changes the locus of the effect from the sensory stage to the decisional stage.

Before the discussing the theoretical and empirical evidence supporting drift rate effects of expectations, it is worth briefly discussing the distinction between *static* and *dynamic* effects of expectations and how they have been mapped to different models in the literature. In this article, we use the term “static” to refer to an effect of expectations that remains constant across the amount of time it takes to complete a single trial. Static effects can be contrasted against “dynamic” effects of expectations, which we use to mean an effect that *changes* over the time it takes to make a choice. The literature often maps the static vs. dynamic distinction onto effects on the starting point vs. drift rate of the accumulation process (e.g., Dunovan et al., 2014; Moran, 2015; Malhotra et al., 2018). This mapping reflects the general interpretation that starting point effects modify the *initial conditions* of the process, whereas drift rate effects modify its *dynamics*. Importantly, however, effects on drift rate can *themselves* be dynamic, such that the rate of accumulation changes over the time it takes to make a single choice (e.g., Afacan-Seref et al., 2018; Hanks et al., 2011). Accordingly, we will use the terms *static drift* and *dynamic drift* to refer to these two different classes of drift rate effects. Finally, our usage of “dynamic” is intended to apply only to effects that occur within a single trial, rather than effects that change over the course of an experimental session (e.g., Alister and Evans, 2026).

#### 2.3.1 Normative grounding for static drift models of expectations

Moran (2015) combined formal decision theory with numerical simulations to investigate how the original DDM can maximize reward rate in an environment that fails to meet the assumptions of SPRT. This was achieved by introducing one simple modification to the formal decision problem: permitting the sensory signal strength to vary across trials. Adding trial-level variability in the sensory signal relaxes the assumption of a homogeneous decision environment (Section 2.2.2.1), which in turn generates uncertainty in the observer’s knowledge of the statistical properties of the environment (Section 2.2.2.2). When an observer does not know in advance how decisive the sensory evidence will be, they cannot appropriately set their threshold in accordance with SPRT, and thus the original DDM fails to provide a normative solution for trading off speed and accuracy in heterogeneous environments (Moran, 2015; Drugowitsch et al., 2012). Despite this, Moran (2015) used numerical simulations to investigate how the process specified by the DDM might approximate optimality (i.e., maximize reward rate) in environments that are both heterogeneous *and* biased (i.e., Π ̸= 0.5). The author found that an additive bias on *both* the starting point *and* drift rate parameters were necessary for maximizing reward rate in this environment, in contrast to previous findings suggesting that no offset on drift was required (van Ravenzwaaij et al., 2012; see Section 4.3.2 of Khoudary et al. (2025b) for an extended discussion of these two findings).

#### 2.3.2 Assumptions and limitations of the normative drift model

The results of Moran (2015) thus indicate that, in biased and heterogeneous environments, a static drift model—coupled with a starting point offset—maximizes reward rate (and thus optimizes the decision process). While these findings advance theoretical knowledge of optimal behavior in more realistic environments, they necessarily rest on simplifying assumptions about the environment and the observer. We discuss each of these assumptions—and their conceptual limitations—before turning to the theoretical framework and computational model we developed to address the gaps left by Moran’s 2015 analysis.

##### 2.3.2.1 Assumption 1: *expectation*-homogeneous decision environment

The first assumption underlying Moran’s (2015) findings is that there is one fixed level of prior probability Π (or bias) in the environment, but that the sensory evidence strength (and, by extension, the drift rate *A*) varies unpredictably across trials. This corresponds to an environment that is *expectation*-homogeneous but *evidence*-heterogeneous: observers use the same expectation for all decisions made in the environment, but the difficulty of the decision varies from trial to trial. We will use the term *fully heterogeneous* to describe environments that have trial-level variability in *both* the sensory signal strength and the prior probability (which we shall denote Π*_cue_* to differentiate from the block-level expectation Π).

As with Assumption 1 for starting point models, basing the normative solution on expectation-homogeneous but evidence-heterogeneous environments restricts the scope of Moran’s (2015) findings to behavior measured in these environments. Experimentally, this corresponds to block-level manipulations of prior probability Π with trial-level manipulations of sensory signal strength. While the lion’s share of early work on expectations in perceptual decisions used exactly this type of decision environment (Carpenter and Williams, 1995; Ratcliff and McKoon, 2008; Simen et al., 2009; Hanks et al., 2011; van Ravenzwaaij et al., 2012), it is becoming increasingly common to measure behavior in fully heterogeneous environments (Kok et al., 2013; Dunovan et al., 2014; Dunovan and Wheeler, 2018; Kelly et al., 2021; Bornstein et al., 2023; Diaz et al., 2024; Khoudary et al., 2025a). To our knowledge, a normative model for this type of behavioral problem has yet to be specified.

##### 2.3.2.2 Assumption 2: perfect knowledge of Π

The second assumption underlying Moran’s (2015) findings is that observers have perfect knowledge of the prior probability Π, but imperfect (or uncertain) knowledge of the signal strength (and corresponding drift rate *A*) on each trial. It is precisely this trial-level uncertainty in sensory signal strength that distinguishes Moran’s findings from the SPRT, and that justifies an effect of expectations on the drift rate in addition to the starting point.

Unlike the assumption of expectation-homogeneity, experimental conditions only sometimes relax this assumption of perfect prior knowledge. Indeed, the dominant approach has been to explicitly instruct humans on prior probabilities, regardless of whether they are manipulated at the block- or trial-level (e.g., Carpenter and Williams, 1995; Ratcliff and McKoon, 2008; Simen et al., 2009; Hanks et al., 2011; Dunovan et al., 2014; Dunovan and Wheeler, 2018; Diaz et al., 2024). In studies using non-human animals, observers are trained on tens of thousands of trials until the experimenters are confident that the animals have sufficiently learned environmental biases (Hanks et al., 2011; Rao et al., 2012), such that there is little to no uncertainty in their knowledge of either Π or Π*_cue_*. As discussed in the context of starting point models (§2.2.2.2), the assumption that observers have near-perfect knowledge of prior probabilities completely elides the questions of *where* expectations come from and *how* the source of expectations might impact their integration with sensory evidence. The theoretical framework and computational model we present in the following sections aim to address this gap directly, positing a central role of *memory retrieval dynamics* in shaping the integration of uncertain expectations with sensory evidence that is itself uncertain.

#### 2.3.3 Empirical support for static drift models of expectations

Before detailing our framework and model, it is worth briefly reviewing some of the recent work supporting static effects of expectations on the drift rate. One of the first studies to report an effect of expectations on drift was conducted by Dunovan et al. (2014), who measured both behavior and neural activity in fully heterogeneous environments where cues unambiguously informed observers about Π*_cue_* on a trial-by-trial basis. The authors found that a “multi-stage model” where both the starting point and drift rate varied as a function of trial-level expectation cue outperformed several simpler models on both quantitative model comparisons and qualitative match to behavioral data. A later study by Dunovan and Wheeler (2018) linked these effects to BOLD activity in category-selective regions of inferotemporal cortex. The authors showed that cue-evoked BOLD timecourses were systematically modulated by the predictiveness of a cue, whereas sensory-evoked timecourses were more strongly modulated by the *mismatch* between a cue’s prediction and the subsequent contents of perception. An EEG study by Rungratsameetaweemana, Itthipuripat et al. (2018)provided corroborating evidence for this “mismatch” effect, which we discuss in detail in Section 5.2.

A study by Kelly et al. (2021) further revealed effects of expectations on drift rate that are modulated by the difficulty of the decision task. Across three blocks, Kelly et al. (2021) manipulated choice difficulty by manipulating the signal strength and amount of time allotted to make each choice. Expectations were manipulated trial-by-trial, with predictive probabilities learned during the practice phase of the task (Kelly et al., 2021). The authors then built a “neurally-informed” sequential sampling model that used EEG measurements to constrain the threshold, non-decision, and starting point terms, and compared the performance of this model to that of the extended DDM (eDDM). In addition to showing that the neurally-informed model outperformed the eDDM on measured and simulated data, the authors reported positive biases on drift rates as a function of expectations across all difficulty blocks (Kelly et al., 2021). Further, these biases were linked to changes in the amplitude of the centroparietal positivity (CPP), an event-related EEG component that has been well-established to co-vary with evidence strength (O’Connell et al., 2012; van Vugt et al., 2019). Specifically, expectation-based effects on the CPP were strongest when observers (1) had a limited amount of time to make a choice and (2) there was a mismatch between the cue’s prediction and the subsequent content of perception (Kelly et al., 2021), further corroborating the findings discussed above.

Finally, a recent study by Liu et al. (2025) showed that effects of expectations on the drift rate and starting point can also be observed in rodents. The authors investigated brain-wide recordings from rodents who learned expectations within a decision environment, and used several formal models to characterize the effects of learned expectations on neural activity. Liu et al. (2025) found that learned expectations modulate the initial value and gain on firing rates both for single neurons and populations thereof, but only if those firing rates appeared to encode accumulated evidence or action initiation rather than sensory evidence. These findings further support the idea that expectations can modify decision—but not sensory-related—processing by modulating both the starting point and drift rate of evidence accumulation on a trial-by-trial basis (Liu et al., 2025).

### 2.4 Section summary

This section reviewed the formal and empirical evidence supporting a *static* effect of expectations on perceptual decisions. Again, our usage of static in this article pertains to *how* the effect of expectations is modeled; a static effect corresponds to an offset on the decision process that does not change over the course of a single decision. On this definition, expectations can—and should—exert static effects on both the starting point and drift rate (Wald and Wolfowitz, 1948; Bogacz et al., 2006; Moran, 2015). We reviewed literature demonstrating behavioral and neural evidence that human and non-human observers optimally incorporate expectations into the decision process, either as a static offset solely on the starting point or on both the starting point and drift rate (Simen et al., 2009; Mulder et al., 2012; Rao et al., 2012; Moran, 2015; Dunovan et al., 2014; Dunovan and Wheeler, 2018; Kelly et al., 2021; Liu et al., 2025).

Mounting evidence, however, suggests that effects of expectations might *vary* within the time it takes to make a choice (Hanks et al., 2011, Rungratsameetaweemana, Itthipuripat et al., 2018, Bornstein et al., 2023, Hachisuka, Shor, Liu et al., 2026). Static starting point models categorically cannot capture such *time-varying* effects of expectations. Static drift rate models can approximate time-varying effects, but are limited in positing effects that are strictly linear in time (Moran, 2015). In the following sections, we suggest—and then show—that time-varying effects of expectations might be driven by the dynamics of *memory retrieval* processes that function to reduce uncertainty during latent state inference processes (Wang et al., 2021; Shadlen and Shohamy, 2016). To do this, we first introduce sequential sampling approaches for modeling the role of memory retrieval in action selection in Section 3.1. Then, we discuss how *uncertainty about expectations* has been omitted from the theoretical study of perceptual decision-making despite its ubiquity in real-world settings, and delineate two dimensions of expectation-uncertainty that have been obscured by assumptions of normative models (Section 3.2; Figure 1). We then present a novel sequential sampling model formalizing the most general construal of our framework (Section 4) and use simulations to show that it reproduces dynamic effects of expectations observed across several different species, tasks, and neuroimaging modalities (Section 5). Finally, we present a secondary analysis of behavioral data to empirically validate a novel prediction generated by our framework of uncertainty-driven memory sampling to further demonstrate its utility for understanding the dynamics of decision-making under uncertainty.

## 3 Conceptual gap: role of memory retrieval dynamics

All mathematical modeling—and normative modeling in particular—requires idealizing properties of the system being modeled (i.e., the *target*; Khoudary et al., 2025b). Idealizations simplify aspects of target systems by *omitting* and/or *distorting*their properties, resulting in formal representations that “selectively attend” to aspects of the system about which model users aim to reason (Portides, 2021). Omitting uncertainty about expectations from formal models—and the experiments used to test them—is a powerful idealization that has led to great progress in understanding the neural and computational mechanisms of optimal decision-making. In doing so, however, the field has systematically ignored the role that *memory retrieval* might play in shaping the dynamics of expectation-guided perceptual decisions.

The relevance of memory for expectation-guided decisions becomes obvious when considering the question of where expectations come from *outside* of laboratory settings; one answer could be previous outcomes of similar decision problems. Because human memory systems store information about past experiences at multiple levels of detail (Tarder-Stoll et al., 2026), precisely what information about the past comes to mind during decision-making—and in what form—is an area of active research (Zeithamova, Schlichting, Preston, 2012; Aronowitz, 2019; Biderman et al., 2020; Wang et al., 2021; Shushruth et al., 2022; Bakkour et al., 2019). For present purposes, we focus on a relatively simple type of information: probabilities of choice outcomes conditioned on a specific context. These “contextual expectations” (Seriès and Seitz, 2013) are acquired through repeated experience with a decision environment (de Lange et al., 2018), and have been shown to facilitate various types of perceptual judgments such as object recognition (Oliva and Torralba, 2007), visual search (Zhou and Geng, 2024), and two-alternative discrimination (Bornstein et al., 2023). Expectation-setting in the real world thus involves retrieving information from memory, but the role of memory retrieval dynamics in shaping perceptual decisions remains largely underexplored. Sequential sampling, or evidence accumulation, provides a common formalism for building theories about how agents trade off speed and accuracy when making decisions based on *both*external (sensory) and internal (memory) information. Combining this formalism with theoretical insights from learning and memory can help advance understanding of the general principles that support flexible and adaptive decision-making in humans, which is what we aim to achieve with the framework and model presented in this article.

### 3.1 Memory as a dynamic source of internal evidence

Retrieving information from memory takes time. Formalizing the relationship between the strength of a memory and the amount of time needed to retrieve it was the goal of the earliest form of the extended drift diffusion model (Ratcliff, 1978), which is now commonly used to analyze behavior in value-based decision-making tasks where the relevant “evidence” used to make a choice is strictly internal to decision-makers (Ratcliff et al., 2016; Krajbich et al., 2010). The *memory sampling* framework explains the ability of accumulator models to fit this class of data by positing that observers sequentially sample information from memory to make predictions about the outcome of each choice, terminating deliberation once the predicted value of one choice sufficiently outweighs the predicted value of the other (Shadlen and Shohamy, 2016; Bakkour et al., 2019; Wang et al., 2021; Biderman et al., 2020; Zylberberg et al., 2024).

Crucially, evidence supporting memory sampling theories of action selection comes from experiments where memory strength is permitted to vary across trials (Wang et al., 2021; Shadlen and Shohamy, 2016; Bakkour et al., 2019; Banavar et al., 2024). This variability derives both from structured manipulations of experimental stimuli (e.g., Nicholas and Mattar, 2026) and the stochasticity of memory retrieval processes both within and across individuals (Wang et al., 2021). Measuring and manipulating memory as a source of decisional evidence has revealed that memory-based decisions obey the same dynamics governing sensory-based decisions: stronger memories lead to faster and more accurate decisions, whereas weaker memories lead to slower and less accurate decisions (Shadlen and Shohamy, 2016; Biderman et al., 2020; Wang et al., 2021).

More recently, it has been shown that humans can adaptively control this memory sampling process, engaging in prolonged sampling only when the expected utility of doing so outweighs the costs of investing more time and effort into the retrieval process (Bornstein et al., 2023; Callaway et al., 2024). The framework of sequential sampling—and its status as a formal approach to partially observable Markov decision problems (Rao, 2010; Drugowitsch et al., 2012; Huang et al., 2012)—permits interpreting this adaptive control of memory retrieval in terms of minimizing uncertainty during latent state inference. When sampling information from an evidence source, observers can take three possible actions: choose latent state +, choose latent state −, or keep sampling. These actions *a* are determined by the distance of the decision variable *x_t_* from the threshold *z* at each point in time as follows:

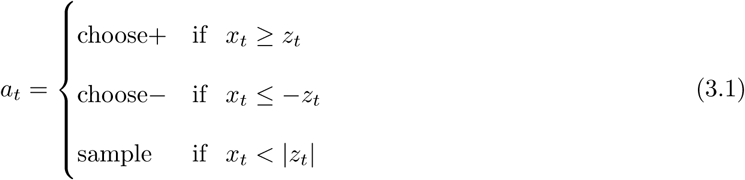

Interpreting the decision variable *x_t_* as reflecting observers’ probabilistic, time-varying belief about the true state of the world (+ or −) further permits interpreting sample actions as decisions to continue acquiring information to reduce uncertainty in *x_t_* (Drugowitsch et al., 2012; Bornstein et al., 2023; Rao, 2010; Kaelbling et al., 1998). Accordingly, prolonged sampling from memory—measured via response times and/or neural activity—can be thought to reflect greater uncertainty about the true value of the data-generating distribution (i.e., the outcome predicted by the set of relevant associative and/or episodic memories). The relationship between sampling time and uncertainty is well-established in the perceptual decision literature, where agents of all species demonstrate robust, systematic relationships between sensory evidence uncertainty and sampling or deliberation time (Gold and Shadlen, 2002, 2007; Hanks et al., 2011; Fetsch et al., 2014; Kiani et al., 2014; Hanks and Summerfield, 2017).

### 3.2 Two dimensions of uncertainty about expectations

Again, a core argument of this article is that the assumptions required for optimal decision processes have obscured the role of uncertainty-driven sampling from memory. To help address this gap, we distinguish two dimensions along which uncertainty about expectations can vary (Figure 1). First, we posit that the *source* of an expectation—or how it is acquired and stored in memory—generates uncertainty about *what* the expectation predicts. Second, we posit that environmental demands on the *deployment* of expectations—or how they are used to guide future decisions—generates uncertainty about *which* expectation to use across consecutive decisions. As shown in the gray text of Figure 1, neural and behavioral evidence for time-varying effects of expectations have been observed in experimental settings corresponding to each of the four quadrants delineated by these dimensions. Section 5 shows that our sequential sampling model formalizing this framework reproduces each of the time-varying effects reported by these studies, suggesting that uncertainty-driven memory sampling offers a unifying explanation for time-varying effects of expectations. In support of this ultimate claim, the rest of this section elaborates upon the dimensions of uncertainty picked out by our framework. For each dimension, we discuss how it relaxes the assumptions of previous normative models, along with the theoretical and empirical motivations for doing so.

#### 3.2.1 Source-related uncertainty

The dimension of source-related uncertainty (orange axis in Figure 1) is intended to capture uncertainty associated with how expectations are acquired and subsequently stored (or represented) in memory. Accordingly, it relaxes the assumptions of perfect knowledge of Π or Π*_cue_* that are required for normative static effects of expectations (Sections 2.2.2.1 and 2.3.2.2). Relaxing this assumption is especially important because it has obscured investigation into *where* expectations come from and *how* the uncertainty associated with an expectation ought to impact its integration with sensory evidence during decision-making.

An observer can be said to have source-related uncertainty about their expectations if they acquire those expectations on the basis of implicit, experience-based learning rather than explicit instruction.^2^ This type of uncertainty is thus about the true prior probability of stimulus occurrence in the decision environment (Π) or the predictive probability of a learned cue with respect to its associated stimulus (Π*_cue_*). A core idea behind our framework is that when prior probabilities are not immediately accessible or directly known by the observer, they must *estimate* these probabilities on the basis of past experience. Doing so requires retrieving relevant information from memory in order to construct an estimate of the prior probability. This process should be jointly modulated by (1) the strength of the true prior probability and (2) the amount of information an observer has stored in memory (or, the precision of their belief). Accordingly, more certain probabilities (closer to 0 or 1) should take fewer samples to estimate, whereas less certain probabilities (closer to 0.5) require prolonged sampling for observers to obtain steady-state estimates. As an observer gains more experience—either with the relevant associations comprising the memory or with estimating probabilities based on a specific set of associations—their belief becomes more precise, and there should be less overall uncertainty in the estimate regardless of its true value. “Structural” expectations—reflecting general statistical information that is preserved across contexts (Seriès and Seitz, 2013)—can be thought to reflect an *asymptotic* form of these expectations learned through experience.

As mentioned above, the majority of experimental work has ignored this dimension of uncertainty about expectations, opting instead to give human observers explicit knowledge of prior probabilities or extensively training non-human animals such that any uncertainty about Π*_cue_* in minimized. One exception to this trend comes from a small handful of experiments where observers implicitly learn environmental biases while performing the main decision task. Behavior and neural activity in both humans (Rungratsameetaweemana, Itthipuripat et al., 2018; Wert et al., 2025) and rodents (Findling et al., 2025) has been shown to be sensitive to these latent environmental biases, with observers of both species demonstrating evidence of learning within as few as 20 trials. Yet how these learned biases manifest in neural representations and the decision process remains underexplored. In humans, for example, these implicitly learned expectations appear to selectively impact decisional, rather than sensory, EEG components (Rungratsameetaweemana, Itthipuripat et al., 2018). Rodent studies, in contrast, have shown that information about the latently-learned environmental bias can be decoded from up to 30% of the entire brain, spanning early sensory to higher-level cortical areas (Findling et al., 2025) rather than being confined to decisional circuitry. These findings thus shed some light on the neural mechanisms involved in integrating uncertain expectations with sensory evidence, but leave open the question of which computations drive this integration. In humans, there is the additional question of *which* representations these computations operate over; the framework and model we develop in this article suggests that this representation is a dynamic, probabilistic “stream” of evidence samples retrieved from associative memory.

#### 3.2.2 Deployment-related uncertainty

The dimension of deployment-related uncertainty (purple axis in Figure 1) captures uncertainty associated with how expectations are cued in advance of decisions. Namely, an observer can be said to have deployment-related uncertainty if they are making decisions in an environment where the prior probability Π*_cue_* differs across decisions in a manner unknown to them. This dimension of uncertainty relaxes the assumption of *expectation-homogeneity* that is ubiquitous among normative models of expectations in decision-making (Edwards, 1965; Bogacz et al., 2006; Simen et al., 2009; Moran, 2015; Malhotra et al., 2018), along with foundational experiments designed to test their predictions (Carpenter and Williams, 1995; Ratcliff and McKoon, 2008; Simen et al., 2009). As shown in Figure 1, deployment-related uncertainty is much lower in experiments that manipulate expectations on the level of blocks of trials, such that observers make hundreds to thousands of consecutive decisions using the same expectation.

Relaxing the assumption of expectation-homogeneity is important for two reasons. First, the assumption itself strongly constrains the space of real-world problems to which existing normative models can be applied. Human observers are rarely tasked with making more than a handful of consecutive perceptual decisions using the same exact expectation, much less hundreds or thousands as in block-level experimental manipulations. Second, and relatedly, it is becoming increasingly common to measure behavior and neural activity of observers making decisions in expectation-heterogeneous environments (Dunovan et al., 2014; Dunovan and Wheeler, 2018; Aitken and Kok, 2022; Kok et al., 2012; Kok et al., 2013; Bornstein et al., 2023; Diaz, Pisauro et al., 2024). Although the prescriptions of existing normative models can offer some guidance for interpreting findings from these environments, a principled model of how deployment-related uncertainty ought to impact decision processes would allow for finer-grained prescriptions guidance for future experimental designs.

### 3.3 Dynamic reliability estimation as a unifying normative principle

A final motivation for relaxing the assumption of expectation-homogeneity comes from the form of previous normative solutions to decision problems in heterogeneous sensory environments. Multiple converging studies have shown that optimal decision-making in evidence-heterogeneous environments—where observers have uncertainty about the strength or duration of sensory evidence—requires some dynamically-changing component of the decision process (Frazier and Yu, 2007; Drugowitsch et al., 2012; Huang et al., 2012; Deneve, 2012; Glaze et al., 2015; Malhotra et al., 2018; Kilpatrick et al., 2019; Barendregt et al., 2022; Callaway et al., 2024; Fang et al., 2026). The intuition behind the need for a dynamic component is the same across all models: when an observer doesn’t know in advance how informative the evidence will be, they must *estimate* its informativeness per unit time—or its *reliability*—over the course of evidence sampling. This evidence reliability estimate is then used to optimize the speed-accuracy tradeoff by dynamically modulating the decision process in real-time. Specifics of how the dynamic adjustment is incorporated vary across model applications and objective functions (see Kilpatrick et al., 2019 for a review), but the general premise of modifying the decision process in proportion to the estimated reliability of evidence remains constant. Our work extends this principle to the integration of expectations into perceptual decisions. We show that multiple disparate empirical observations of time-varying effects of expectations can be explained by a single model that (1) treats expectations as a dynamic “stream” of samples from memory and (2) dynamically adjusts their weighting in the decision process according to a time-evolving belief about the reliability of sensory evidence *relative* to that of the expectation.

## 4 Dynamic reliability-weighted multi-source sequential sampling model

This section presents a sequential sampling model formalizing the conceptual framework developed above. One of its core features is modeling expectations as an independent source of probabilistic information that observers sample from and accumulate *in parallel* to sensory evidence. This process thus equips observers with two sources of probabilistic evidence that they have to combine to make decisions. We draw on well-established theoretical and empirical findings in multisensory integration to propose that these two evidence sources—memory and sensation—are combined according to their *relative* reliability (e.g., Fetsch et al., 2011; Landy et al., 2011; Angelaki et al., 2009). We model the reliability estimation process as dynamically updating with each new sample of noisy evidence from each source (Figure 2); this eliminates the need for using elapsed time as a proxy variable, and encompasses cases where proxies such as time diverge from evidence reliability. The following section shows how this dynamic, relative reliability-weighted process captures time-varying effects of expectations observed by Hanks et al. (2011), Rungratsameetaweemana, Itthipuripat et al. (2018), Hachisuka, Shor, Liu et al. (2026), and Bornstein et al. (2023).

**Figure 2:**
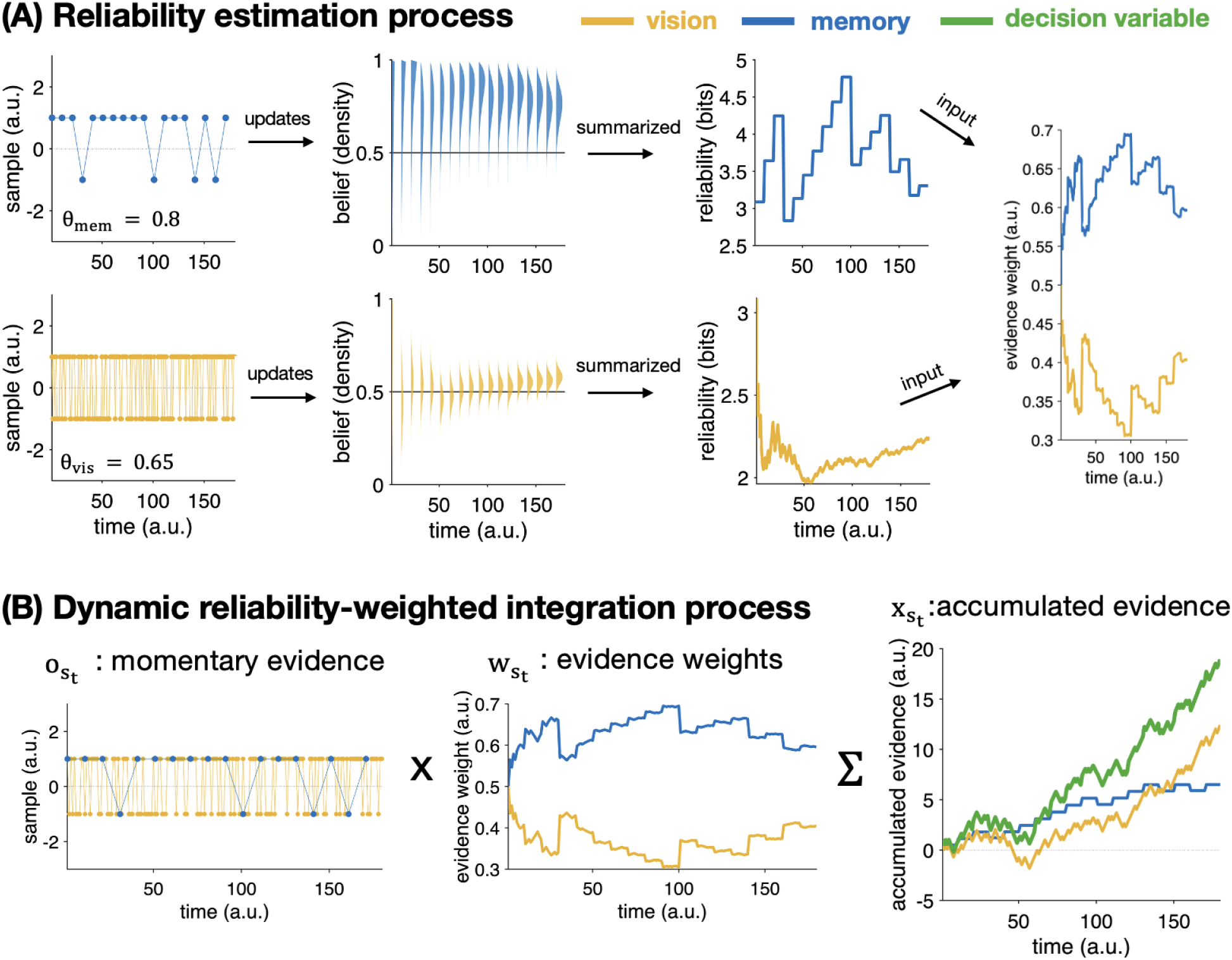
Graphical depiction of single-trial dynamics in our model. Quantities corresponding to memory are plotted in blue, quantities corresponding to vision are plotted in yellow, and quantities corresponding the decision variable are plotted in green. Figures were generated using a vision sampling rate of 60Hz, a memory “sampling rate” of 6Hz (*γ* = 10), and an encoding noise ratio *ɛ* = 0.1. (A) Model components involved in the dynamic reliability estimation process specified by our model. The momentary evidence *o_s_* is generated by a Bernoulli distribution with *p* = *θ_s_* and assumed to be corrupted by Gaussian noise (Equation 4.1). The true value of *θ_s_* defines how informative each evidence source *s* is with respect to the two-alternative choice. Each new sample *o_s_* updates a Beta-distributed belief *g_s_* about the value of *θ*. We assume that the observer’s internal estimate of each source’s reliability *λ_s_* is equivalent to the inverse of the dot product of *g_st_* with the information entropy function. The relative magnitude of each of these reliability estimates *λ_s_* then defines how an observer weights each source of evidence at each moment in time (*w_s_*). (B) Model components depicting the reliability-weighted evidence integration process. We assume that observers weight each evidence source by the dynamic relative reliability estimate computed above, and sum these samples over time to create a decision variable (green) that is a reliability-weighted blend of memory (blue) and sensory (yellow) evidence.

### 4.1 Model specification

In our model, simulated agents sequentially sample evidence from two sources *s* = {*memory, vision*}. Samples for both sources are generated by a Bernoulli distribution with parameter *θ_s_*, which we assume to be unknown by the observer (Figure 2A, left). Additionally, we assume that these samples are noisily encoded at a proportion defined by *ɛ*, such that:

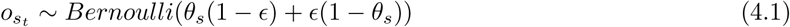

Next, we posit that observers use these samples to form and update a Beta-distributed belief *g_s_*(*t*) about the value of *θ_s_* for each evidence source (Figure 2, middle). On this specification, *θ* corresponds to the “signal strength” for each evidence source: the true *θ_vis_* value is identical to signal strength *A* for sensory evidence, and the true *θ_mem_* value is identical to predictive probability Π*_cue_*. Each new sample from each source updates the belief *g_s_*(*t*) according to:

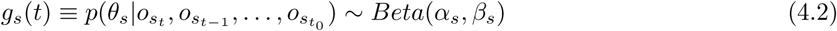

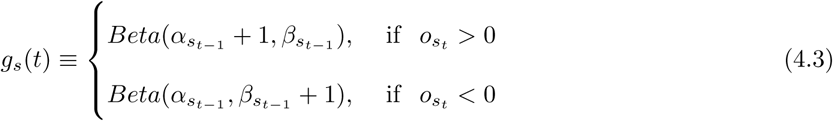

The central tendency of *g_s_*(*t*) captures the observer’s posterior belief about each source’s likely signal strength at each moment in time, and the variance of *g_s_*(*t*) captures the observer’s uncertainty about the inferred signal strength. We define internal evidence reliability *λ_st_* as a term that captures both the central tendency and variance of *g_s_*(*t*) (Figure 2A, right):

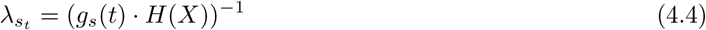

where X is a vector of probability values ranging from 0 to 1 and H is the information entropy function. Each source’s dynamic reliability estimate thus corresponds to the dot product of the observer’s belief about that source’s signal strength *θ_s_* with the information entropy function *H*(*X*). This specification creates a reliability estimate that quantifies how informative an ideal observer ought to believe a particular evidence source is, based on how informative it has been up to this point in the trial. Then, we define the optimal weighting of each evidence source *w_st_* using the standard ideal-observer model of cue integration (Figure 2B, left):

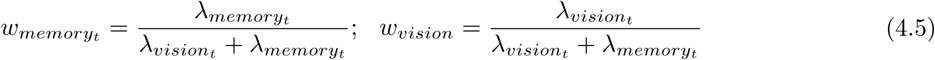

Observers weight each observation from each evidence source by these source-specific weights, creating a decision variable *x_t_* that is a time-varying reliability-weighted sum of samples from independent streams of memory and visual evidence (Figure 2B, right):

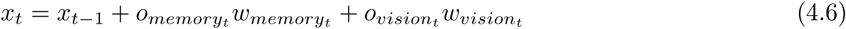

The decision process terminates according to

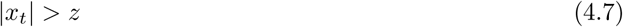

where *z* is a scalar defining the boundary separation (threshold) value.

In addition to the integration of *x_t_* over time, our model posits independent reliability-weighted accumulators for memory and vision that represent the unimodal evidence sampled from each source (Figure 2B, right):

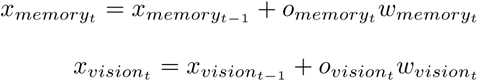

Additionally, the model assumes that memory samples are generated at a fraction of the rate that visual samples are received by the observer, such that the total number of samples generated by each evidence source (memory and vision) is given by:

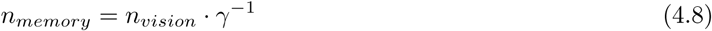

where *γ* is a scalar. All findings in this article assume that *γ* scales the “rate” of memory evidence generation such that memory samples are delivered along the range of theta oscillatory frequencies that have been associated with task-induced memory retrieval (2-12Hz; Jacobs, 2014; Kragel et al., 2020; Vivekananda et al., 2021; Seger et al., 2023). For the majority of simulations, this corresponded to *γ* ∈ {30, 20, 15, 12, 10, 8, 7, 6, 5}.

### 4.2 Steady-state model dynamics

To further elucidate the dynamics generated by our model, we display steady-state timeseries of the relative evidence weights generated across different combinations of memory and sensory evidence strength (Figure 3). Steady-state timeseries were obtained by averaging across 5,000 simulations of 100 decision trials for each combination of memory evidence strength *θ_mem_*, sensory evidence strength *θ_vis_*, encoding noise *ɛ*, and memory “sampling rate” *γ*. We simulated the process with *θ_s_* ∈ {0.5, 0.65, 0.8}, *ɛ* ∈ {0.01, 0.06, 0.11, 0.15, 0.2}, and *γ* ∈ {30, 20, 15, 12, 10, 8, 7, 6, 5, 1}, and *z* = ∞. For simulations where *θ_mem_ >* 0.5, we included only the subset of trials where memory and sensory evidence were in agreement about the correct decision (i.e., cues expectations were “valid” or “congruent” with sensory evidence).

**Figure 3:**
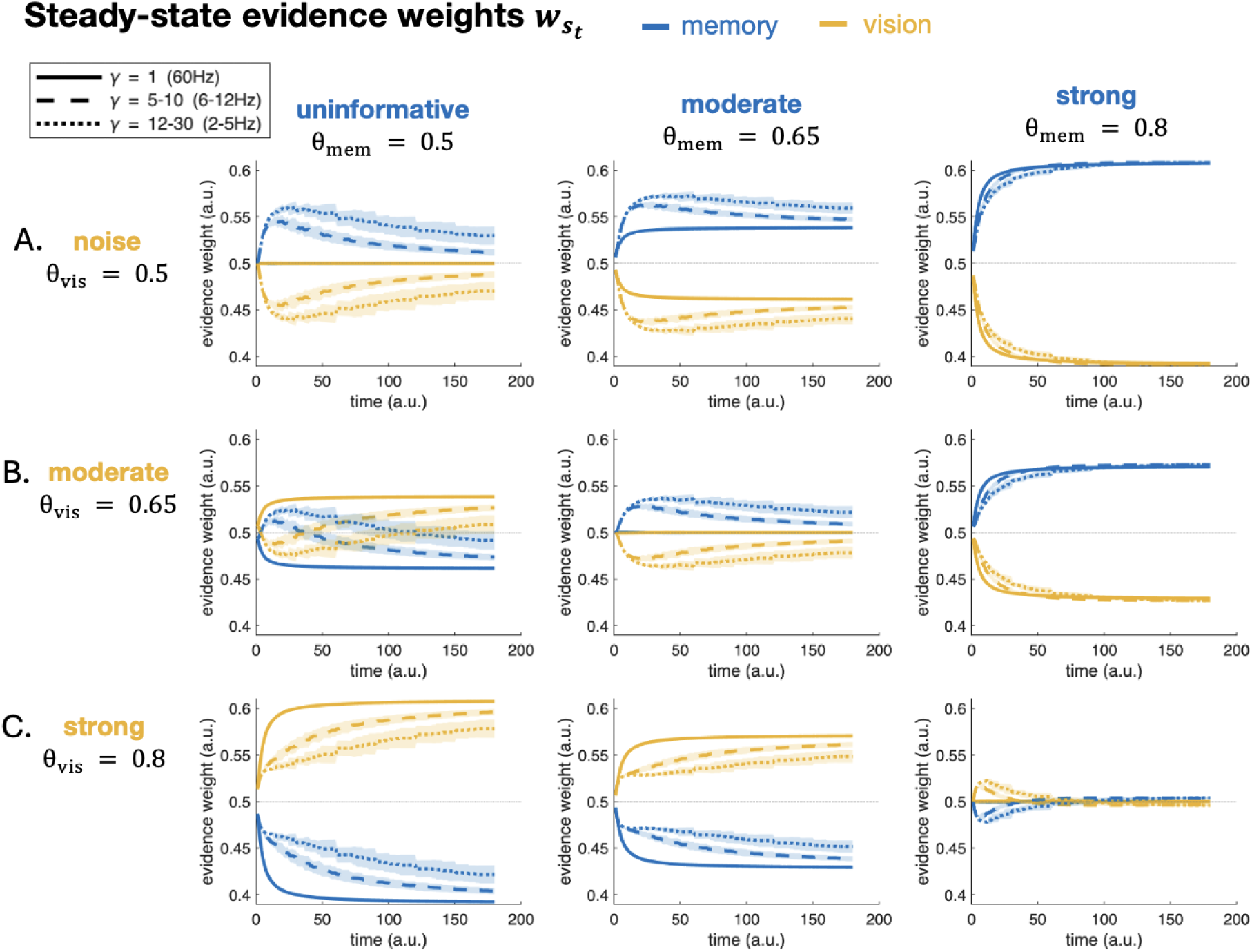
Steady-state dynamic weights on memory (blue) and sensory (yellow) evidence. The rate at which memory evidence enters the decision process is indicated by different line styles (solid = equal rate as vision, dashed = high theta frequencies, dotted = low theta frequencies). (A) Weights when sensory evidence is totally uninformative (*θ_vis_* = 0.5). (B) Weights when sensory evidence is moderately informative (*θ_vis_* = 0.65). (C) Weights when sensory evidence is strongly informative (*θ_vis_* = 0.8). Shaded regions correspond to standard error computed across all levels of *ɛ* and the subset of *γ* corresponding to each linetype.

Figure 3 demonstrates how the steady-state timeseries change across different combinations of *θ_s_* and *γ*. Importantly, we display timeseries with *γ* = 1—corresponding to identical memory and vision “sampling rates”—in solid lines to aid in interpreting the effects of *γ* on our process. These timeseries, along the diagonal of Figure 3 where *θ_vis_* = *θ_mem_*, validate that there ought to be equal weight on both evidence sources when they are equally informative *and* enter the decision process at the same rate. The dashed and dotted lines on the diagonal plots, however, show that slowing the rate of memory evidence sampling relative to vision generates a transient *upweighting* in one of the sources relative to the other. Specifically, for sufficiently weak sensory evidence (*θ_vis_ <* 0.8), expectations are upweighted early on relative to vision, with the degree of upweighting scaling inversely with *θ_vis_* (top left and middle panels of Figure 3). When sensory evidence is sufficiently strong, however, *vision*is transiently upweighted before the quickly collapse to being identical (bottom right panel of Figure 3). Further, the degree to which equally-informative expectations are upweighted relative to vision scales with how quickly memory samples become available for the observer: memory is more strongly upweighted for evidence generated at the low theta frequency (dotted lines) relative to the high theta frequency (dashed lines).

Panels *off* the diagonal in Figure 3 further display dynamics when memory and sensory evidence differ in their informativeness. The top row of Figure 3 shows that, when sensory evidence is completely uninformative, memory remains consistently upweighted throughout the decision process. Comparing across moderate (top row, middle) and strong (top row, right) expectations shows that both the magnitude and rate at which memory is upweighted scales as a function of its informativeness. The middle row of Figure 3 shows that moderately strong sensory evidence elicits heterogeneous dynamics across differently-informative expectations. When expectations carry no information (middle row, left) *and* they become available at a slower rate than sensory information (dashed and dotted lines), they are again transiently upweighted relative to sensory information. However, as the reliability estimates *λ_s_* converge on their steady-state values over the course of sampling, vision becomes *upweighted* relative to memory. And again, the *rate* with which this “crossover” occurs depends on how quickly memory evidence becomes available to the observer (dashed vs. dotted lines). When expectations are *more* informative than sensory evidence, however, memory is quickly and persistently upweighted relative to vision (right panel in Figure 3B). Finally, the bottom row of Figure 3 shows that when sensory evidence is sufficiently strong (*θ_vis_*), it *always* ought to be weighted more than memory evidence that is uninformative (left panel of Figure 3C), moderately informative (middle panel of Figure 3C), or even equally informative but arriving at a slower rate (right panel of Figure 3, dotted and dashed lines).

Taken together, the dynamics in Figure 3 capture core theoretical ideas in our model. When an evidence source has continuously provided reliable information throughout a decision (i.e., when *θ_s_* is large), that source should increasingly bias one’s decision over time. However, when an evidence source has continuously provided weak or unreliable information throughout a decision (i.e., when *θ_s_* is small or null), the *alternative* evidence source should increasingly be relied on over time. That these effects are modulated by the relative *rate* at which samples become available to the observer further underscores the importance of considering the temporally-extended nature of memory retrieval, and how those dynamics shape the integration of uncertain expectations with sensory evidence that is itself uncertain.

## 5 Dynamic reliability-weighted integration unifies time-varying effects of expectations

This section simulates the model specified above under four different experimental conditions, each corresponding to a unique quadrant in Figure 1. In doing so, we demonstrate that our conceptual framework of uncertainty-based memory sampling—and the dynamic reliability-weighted model it inspired—offers a unifying explanation for time-varying effects of expectations on perceptual decisions. Crucially, each of the findings simulated below come from different research groups, experimental designs, and neuroimaging modalities (local field potentials, EEG, fMRI, and intracranial recordings), and were accordingly explained by different neurocomputational processes. The ability of our model to capture latent dynamics across this broad range of explanatory targets suggests that it identifies a core computational process driving the integration of prior knowledge with incoming sensory information.

All simulations were carried out in MATLAB 2023B and 2025A using scripts available on Github. Details about each simulation are contained in its corresponding subsection below.

### 5.1 Quadrant I: block-level instructed expectations (Hanks et al., 2011)

The first quadrant of Figure 1 corresponds to instances when we predict time-varying effects of expectations to be most subtle: expectations come from a source with high certainty (direct instruction) and there is no uncertainty about which expectation to use across decisions (block-level deployment). Intriguingly, however, one of the first and most influential observations of time-varying effects of expectations were reported from a task using exactly these settings (Hanks et al., 2011).

#### 5.1.1 Study design & key findings

Hanks et al. (2011) measured behavior and local field potentials from humans and macaques performing a standard motion discrimination task. Humans were explicitly instructed about the block-level prior probabilities, and macaques were extensively trained until their behavior demonstrated successful learning of block-level reward probabilities, effectively equipping them with instructed expectations. Sensory evidence coherence, or signal strength, was allowed to vary across trials, generating an *evidence-heterogeneous* but *expectation-homogeneous* decision environment in each block. With this design, Hanks et al. (2011) reported behavioral, neural, and computational evidence indicating that observers nonlinearly increase their weighting of the prior probability Π as a function of elapsed decision time (Figure 4A, left). Further, this effect was greatest on trials where sensory evidence was maximally uncertain (Hanks et al., 2011).

**Figure 4:**
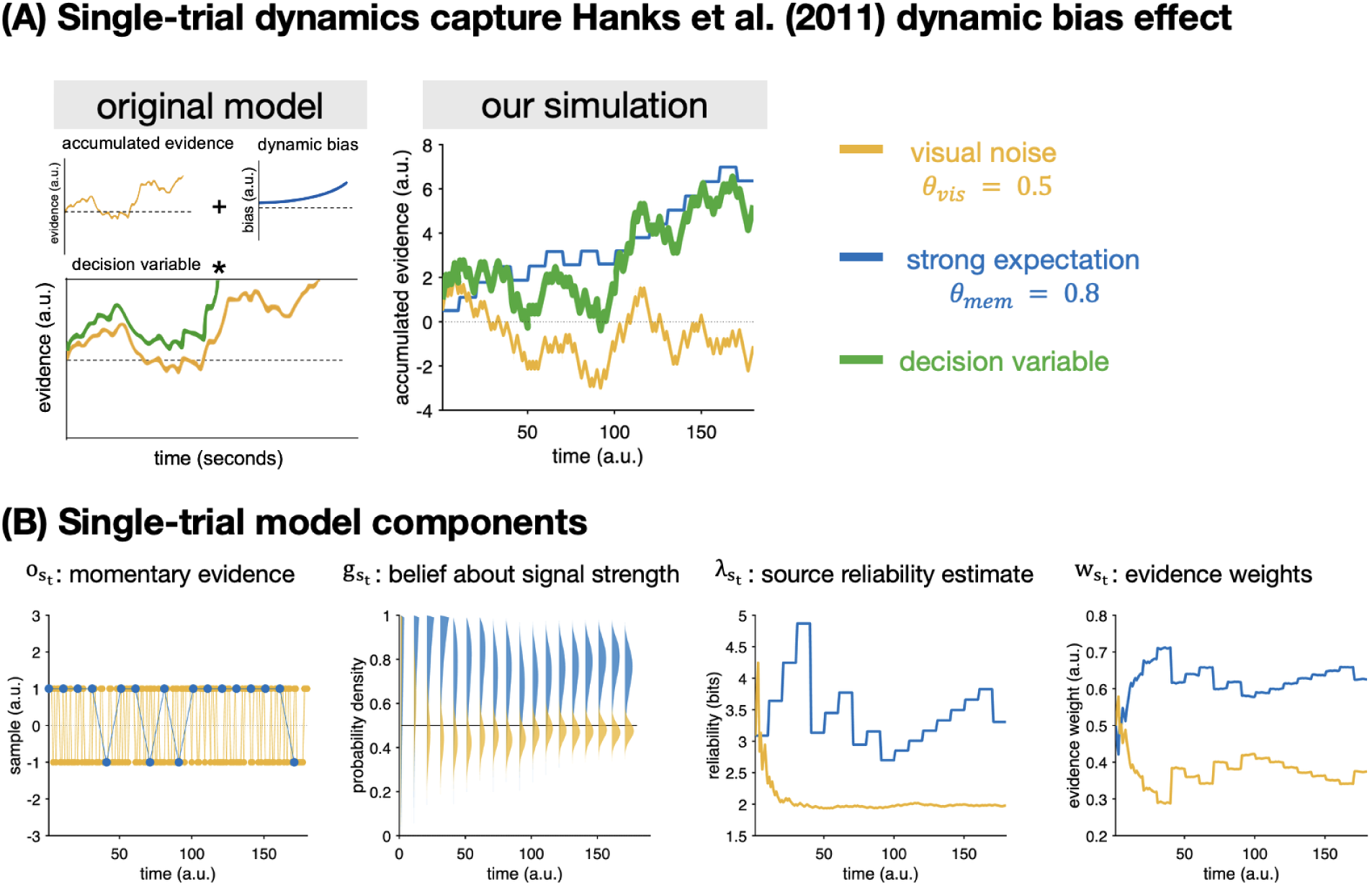
Dynamic reliability-weighted integration captures Hanks et al. (2011) dynamic effect of expectations. In all figures, memory evidence (expectations; Π) is plotted in blue, vision evidence is plotted in yellow, and the decision variable is plotted in green. (A) On the left, a stylized reproduction of the original model used by Hanks et al. (2011) to explain their “late effect” of expectations. On the right, a representative trial from our model demonstrating the same single-trial dynamics. (B) Model components involved in generating our representative trial above.

#### 5.1.2 Computational modeling

The authors explained these findings by theorizing that observers learn a *time-dependent accuracy* (TDA) function as they make decisions in evidence-heterogeneous environments. Learning such a function is behaviorally advantageous because it allows observers to optimize their speed-accuracy tradeoffs without having full knowledge of the statistics of the environment. Instead, they can leverage the systematic—and even idiosyncratic—association between sensory evidence strength and the amount of time needed to commit to a decision to *infer* signal strength *A* based on elapsed time (Hanks et al., 2011). A *dynamic bias signal* (DBS) can then be defined based on the ratio of the prior probability to time-dependent accuracy. Formally,

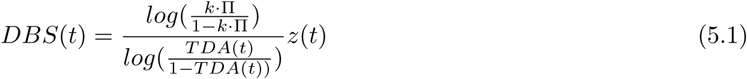

where Π is the prior probability of a motion direction, *z* is the height of decision boundary, and *k* is a constant that scales Π to accommodate cases where primate observers misestimate the true prior probability. The time-dependent accuracy function (TDA) was computed based on observers’ behavior in the unbiased blocks, and then was used to fit behavior in the biased blocks (Hanks et al., 2011).

Although not explicitly described as such, Hanks et al.’s (2011) dynamic bias signal implements a dynamic reliability-weighted process very similar to the one we specify in Section 4.1. Rather than estimate evidence reliability directly, however, Hanks et al., 2011 suggest that evidence reliability is inferred based on elapsed decision time. The inference about evidence reliability is thus given by the association of longer deliberation time with lower coherence stimuli, which also generally are associated with a lower degree of choice accuracy. However, as we will show next, these same dynamics can be generated by our reliability-weighted process which does not assume or require that observers learn time-dependent accuracy functions.

#### 5.1.3 Simulation methods

The goal of our simulations was to show that our relative reliability-weighted process generates the same “late effect” observed by Hanks et al. (2011): greater weighting of expectations *late* in the decision process, especially when sensory evidence is weakly informative. To do so, we generated timeseries data using *θ_vis_* = 0.50, *θ_mem_* = 0.8, *ɛ* = 0.11, and *γ* = 10 with a visual presentation rate of 60 Hz (effective memory “sampling rate” = 6Hz). These parameter settings correspond to the experimental conditions where the late effect was most prominent (Hanks et al., 2011). We discretized the continuous-time simulation such that each timestep corresponded to one “frame” of presented visual evidence; each trial’s three second simulation thus consisted of *n_vision_* = 180 and *n_memory_*= 18. Importantly, our explanation for the effects observed by Hanks et al. (2011) (Figure 4) are independent of threshold *z*, which was set at an arbitrary value of *z* = 250.

#### 5.1.4 Simulation results

Figure 4A demonstrates that, at a single-trial level, our model (right) produces the same dynamics as Hanks et al.’s (2011) dynamic bias model (left). Crucially, however, our model suggests that the “dynamic bias signal” is comprised of an independent *memory accumulator* that samples and integrates evidence generated by memory in parallel to the sensory evidence presented by the observer.

Figure 4B displays the corresponding model components for this representative trial. The model components generating these dynamics are displayed in (Figure 4B). Just as each sample of visual evidence *o_visiont_* contributes to the observer’s estimate of visual evidence reliability *g_visiont_*, so too does each “sample” of memory evidence *o_memoryt_* update the observer’s estimate of memory evidence reliability *g_memoryt_*. Because the signal strength of vision is so low relative to the evidence provided by memory, the expectation is continuously estimated as being more reliable (or informative) throughout the trial, leading to consistently higher weights on memory than sensory evidence (Figure 4B, rightmost plots). When these dynamic, relative-reliability defined weights are used to integrate the sensory and memory evidence over time, they generate the timeseries plotted in Figure 4A (right), where the decision variable becomes increasingly biased toward the expectation over the course of a single decision. Further, the rightmost panel in Figure **??**A further demonstrates that the increased weighting over time is present in the steady-state dynamics of our model averaged across several plausible values of *γ* and *z*.

Taken together, these simulation findings demonstrate that the relative reliability-weighted process we specified to account for the role of uncertainty about expectations can capture a canonical time-varying effect of expectations: increased weighting as a function of elapsed decision time (Hanks et al., 2011). Crucially, these effects were observed in experimental settings where we predict uncertainty-driven time-varying effects to be the most subtle (Quadrant I of Figure 1), and thus serve as a strong test of the plausibility of the framework and model that we propose. The following simulation findings further demonstrate the ability of our model to capture time-varying effects observed in even more uncertain circumstances.

### 5.2 Quadrant II: block-level learned expectations (Rungratsameetaweemana, Itthipuripat et al., 2018)

An active debate in the perceptual decision-making literature is whether expectations can modulate sensory responses themselves (Kok et al., 2012; Kok et al., 2013; Afacan-Seref et al., 2018) or only “post-sensory” decision processes (Summerfield and de Lange, 2014; Rungratsameetaweemana and Serences, 2019). Specifically, it has been argued that trial-level expectation cues can elicit *attentional* effects on sensory responses that are conceptually distinct from those driven by prior knowledge of stimulus probabilities (Summerfield and de Lange, 2014). To address this question, Rungratsameetaweemana, Itthipuripat et al. (2018)designed an experiment where observers had to learn block-level expectations *during* a perceptual decision task. This in-context learning of prior probabilities bypasses the concern about attentional effects elicited by cue-based manipulations of expectations, such that any effects on sensory responses must be attributed to learned expectations about prior probabilities.

#### 5.2.1 Study design & key findings

Rungratsameetaweemana, Itthipuripat et al. (2018)report behavioral and EEG data from n=17 human subjects performing the task. Sensory stimuli were comprised of small red and blue bars whose locations and orientations were refreshed at slow (33.33 Hz) and fast (50 Hz) refresh rates. Across all decisions, the signal-to-noise ratio of sensory evidence was fixed at 0.685. The authors independently manipulated expectations about color, orientation, and motor response across blocks comprised of 60 trials each, but fixed the magnitude of the prior probability to Π = 0.7 across all blocks. This means that in biased blocks, *n* = 42 trials were presented with the dominant stimulus feature whereas *n* = 22 trials were presented with the other stimulus feature. Crucially, neutral blocks presented both stimulus features equally, and subjects completed 4 blocks of 60 trials under neutral expectations as well.

With this design, the authors showed that expectations can exert robust effects on choice behavior in the absence of modulations to sensory processing as measured by EEG. Specifically, Rungratsameetaweemana, Itthipuripat et al. (2018)showed that expectations had no effect on the early visual negative potential, an event-related EEG component that indexes sensory information processing. However, the authors did observe effects of expectations on the centroparietal positivity (CPP), a different event-related EEG component that is strongly associated with the decision variable in accumulator models (O’Connell et al., 2018), as well as effects of expectations on the amplitude of parietal alpha and frontal theta measurements. Crucially, all of these effects occurred *after* each component reached its peak value for the decision, *and* were driven by trials where the sensory evidence was *unexpected* (or *incongruent* with the learned expectation for that block). In what follows, we show that our dynamic reliability-weighted integration process can capture the dynamics observed in frontal theta amplitude.

#### 5.2.2 Simulation methods

Simulations were run with *θ_vis_* = 0.685, *θ_mem_*= 0.7 for biased blocks, and *θ_mem_* = 0.5 for neutral blocks. Although the signal-to-noise ratio of sensory evidence was held constant across trials, the authors manipulated the rate of sensory evidence presentation randomly across trials. Accordingly, we ran simulations with both the low (33.33 Hz) and high (50 Hz) flicker rates used in the original study. This simulation’s aim of capturing neural activity in the theta frequency further aided in constraining the range of *γ* values used in the low and high flicker rate conditions. We defined the range of *γ* values such that memory evidence was generated across the range of theta activity observed in humans (1.5-8Hz; Rungratsameetaweemana, Itthipuripat et al., 2018; Jacobs, 2014). Simulations with low flicker rates used *γ* ∈ {4, 6, 8, 10, 12, 20} corresponding to activity at 8.25, 5.5, 4.125, 3.3, 2.5, and 1.5Hz, and simulations with high flicker rates used *γ* ∈ {4, 6, 8, 10, 12, 20} to generate memory samples at 8.3, 6.25, 4.16, 3.5, 2.5, and 1.5Hz, respectively.

With these free parameters of the model constrained by the original study, we then defined the range of threshold values *z*∈ {3, 4, 5} based on the similarity between trial-averaged behavioral data produced by the model in simulation and by human observers in the experiment. Importantly, this trial-averaged behavior was generated by simulated *n* = 18 “subjects”, each with a unique combination of memory sampling rate *γ* and threshold *z*. Each “subject” contributed the same number of trials as completed by human subjects in the experiment: 360 biased trials and 120 neutral trials at each flicker rate (720 biased and 240 neutral total). We additionally varied evidence encoding noise *ɛ* at the whole-sample level, such that each run used *ɛ* ∈ {0.01, 0.06, 0.11, 0.15, 0.2}. Figure 5 displays results generated with *ɛ* = 0.11, and Figure **??**A displays results averaged across all values of *ɛ*. As in the original experiment, the first 1000ms of each trial were comprised of sensory noise (*θ_vis_* = 0.5). Then, sensory evidence was presented for 850ms at either the low or high flicker rate, before being replaced by sensory noise for 600ms (Rungratsameetaweemana, Itthipuripat et al., 2018). Each timestep in the simulation corresponded to one frame of evidence, such that slow flicker trials had *n* = 82 timesteps and fast flicker trials had *n* = 124 timesteps.

**Figure 5:**
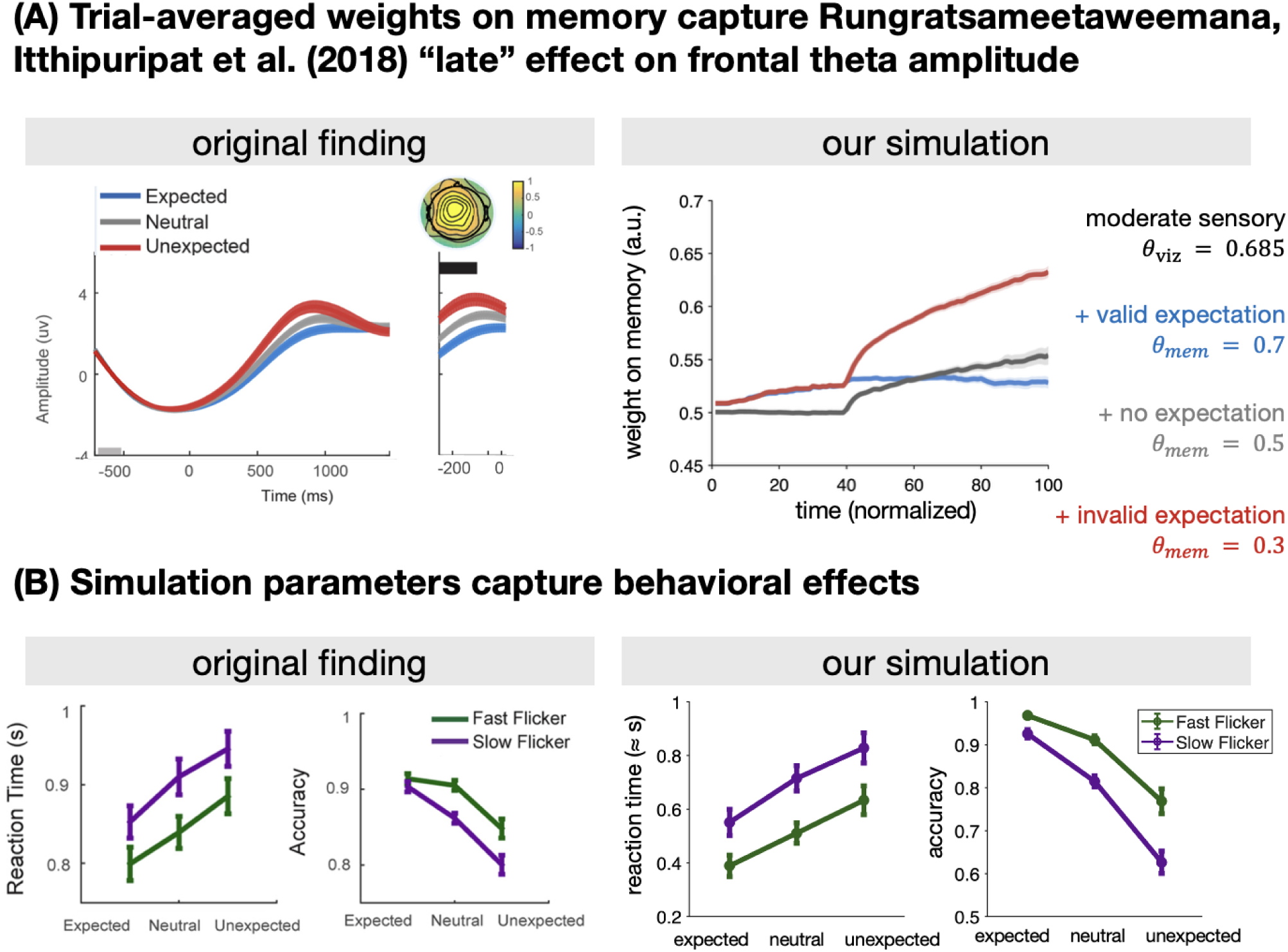
Dynamic reliability-weighted integration captures time-varying effect of expectations observed by Rungratsameetaweemana, Itthipuripat et al. (2018). (A) Original finding, left: violations of expectations increase frontal theta amplitude at later timepoints in the trial. Our simulation, right: weights on memory— interpretable as an incentive to sample evidence from memory—are greater for unexpected stimuli (red) relative to neutral (gray) and expected (blue). Crucially, the effect increases over time and is greatest at the end of the decision. Shaded areas reflect standard error across 15 simulated subjects. (B) Original finding, left: flicker rate and expectation manipulations induce canonical modulations of accumulation-based choice behavior. Slow flicker trials (purple), where fewer samples of visual evidence are presented per unit time, lead to slower reaction times and lower choice accuracy relative to fast flicker trials (green). Our simulation, right: choices and reaction times generated from our model replicate these behavioral effects, validating our choices for sensory signal strength *θ_vis_* and threshold *z*. Errorbars in both (A) and (B) reflect standard error across *n* = 18 simulated “subjects”, each of which had a unique value of *γ* and *z*.

The timeseries plotted in Figure 5A (right) used a normalization procedure to align memory weights generated in the low and high flicker simulations onto a common numerical space. For each subject, weights were averaged across trials in each of the expectation x flicker rate conditions, and then interpolated onto a common grid comprised of n=100 timesteps. Once normalized, weights were then averaged across flicker levels within each participant, and then averaged across participants to obtain condition-specific group means and standard errors. Group-averaged timeseries for each of the low and high flicker conditions in their native numeric spaces are displayed in Figure **??**B.

#### 5.2.3 Simulation results

Figure 5A shows that our model’s time-evolving weights on memory capture the “late” effect that Rungratsameetaweemana, Itthipuripat et al. (2018)observed on frontal theta amplitude. After the initial period of sensory noise, weights on memory slowly increase over time in all three expectation conditions. However, the rate of increase is largest on trials when the expectation is *invalid* with respect to sensory evidence (red line), relative to trials where the expectation carries no information about sensory evidence (gray line) or valid/congruent information about sensory evidence (blue line).

This effect emerges naturally from positing that observers use the same time-constant boundary to terminate their decisions across different experimental conditions (low/high flicker, expected/neutral/unexpected). Because evidence weights in our model are estimated dynamically over the course of sampling, trials where observers take longer to make a choice naturally generate weights on memory that continue to increase relative to trials where observers terminate their decisions earlier. Crucially, this only holds if the expectation magnitude *θ_mem_* and sensory evidence *θ_vis_* are held constant across trials, which is precisely the case when comparing valid and invalid trials. Neutral trials appropriately generate weights whose magnitudes are intermediate between valid and invalid. However, because there is no true information in the neutral expectation signal, its timeseries is quantitatively indistinguishable from valid expectations that don’t need to be sampled because a choice has already been made.

Figure 5B additionally shows that the ranges of *γ* and *z* used in our simulation capture the key behavioral effects, validating the link between our model’s latent processes and the EEG measurements in the original study. Crucially, in doing so, we demonstrate that our model can reproduce both behavior and neural dynamics in environments corresponding to Quadrant II in Figure 1: learned expectations manipulated on the level of blocks. Our findings further raise an alternative interpretation of the frontal theta effects observed by Rungratsameetaweemana, Itthipuripat et al. (2018): rather than reflecting executive control processes, the late increase in theta amplitude may reflect prolonged sampling of learned expectations on unexpected trials relative to neutral and expected.

### 5.3 Interim summary

Taken together, Sections 5.1 and 5.2 show that dynamic, relative reliability-weighted integration of independent memory and sensory accumulators can reproduce dynamic effects of expectations observed in multi-modal, cross-species neural measurements. Both of these simulations focused on dynamics observed in *expectation-homogeneous* environments, where observers have no uncertainty about which expectation to use across decisions. The key factor differentiating the two simulations is whether those expectations were acquired through instruction (Section 5.1) or learned through experience with the decision environment (Section 5.2). Both types of block-level expectations exerted time-varying effects that occurred *late* in the decision process: with the Hanks et al. (2011) increasing nonlinearly over elapsed time, and the Rungratsameetaweemana, Itthipuripat et al. (2018)effect occurring after the peak of the ERP signal on each trial. We turn next to demonstrating that our model can capture existing findings corresponding to the remaining two quadrants in Figure 1, where observers have uncertainty about *which* expectation to use for each decision.

### 5.4 Quadrant III: trial-level learned expectations (Bornstein et al., 2023)

On our framework, experimental settings corresponding to Quadrant III are those where we expect time-varying effects of expectations to be most pronounced: there is source-related uncertainty about *what* the true value of the expectation is, as well as deployment-related uncertainty about *which* expectation will be used on every trial. A recent study by Bornstein et al. (2023) measured human behavior and neural activity in precisely these settings and found evidence for uncertainty-mediated time-varying effects of expectations that occur *early* in the decision process, before any sensory evidence has been presented.

#### 5.4.1 Study design & key findings

Bornstein et al. (2023) utilized a two-phase task design, where observers first learn probabilistic cue-stimulus relationships in a learning phase, and then make cue-guided perceptual decisions in a decision-making phase. The expectation cues consisted of colored fractals, and the perceptual stimuli were grayscale scene and face images. The decision task consisted of two-alternative discrimination between two images from the same category, e.g., reporting whether the dominant stimulus on each trial was Face A or Face B. Accordingly, observers learned that each fractal makes a unique prediction about which image is more likely. Crucially, images from the same category (faces vs. scenes) were equally predicted by their corresponding fractal, such that, for example, faces were strongly predicted by their fractals (Π*_cue_* = 0.8) whereas scenes were weakly predicted by their fractals (Π*_cue_* = 0.6). These learning conditions thus created both item- and category-specific predictions, such that, for example, the correct answer for face decisions was more predictable than the correct answer for scene decisions. Finally, during learning, participants only had a small amount of exposure to each cue-stimulus pairing (∼ 30 trials) to ensure that there was learning-related uncertainty around each expectation.

Sensory evidence during the decision-making phase was presented at an effective rate of 30Hz: frames were updated at 60Hz, but each frame of signal was forward- and backward-masked by noise (phase-scrambled superpositions of the to-be-discriminated images). The difficulty of the decision was manipulated by varying the proportion of frames containing the target image, such that trials with closer to equal frames of target and lure images were more difficult to resolve. Each trial in the decision phase began with a brief (750ms) presentation of a fractal image followed by an interstimulus interval that varied uniformly across 4000, 6000, and 8000ms. Once the ISI elapsed, sensory evidence appeared on screen and observers had up to 3000ms to make their response. Crucially, the authors yoked decision difficulty to the category membership of the sensory evidence; e.g., in a given block, face evidence would always have a low signal-to-noise ratio whereas scene evidence would always have a high signal-to-noise ratio. Combined with the category-level expectations created during learning, this manipulation created *joint* about sensory evidence on each trial: a probabilistic expectation about the correct answer (defined by Π*_cue_*) and a *deterministic* expectation about the difficulty of the upcoming decision (learned through experience in the decision phase).

Because each cue gives observers deterministic information about the relative signal strength of the upcoming decision, this design allowed Bornstein et al. (2023) to identify behavioral effects in line with predictions of normative starting point models. Specifically, on as many as a third of trials with a high Π*_cue_*that also predicted a low upcoming signal strength *A*, observers opted to make a response *before* observing any sensory evidence at all. This prediction follows directly from normative starting point models, which show that as Π approaches 1 and decisions approach maximum difficulty, the optimal procedure is to respond immediately on the basis of the expectation (Simen et al., 2009).

An important goal of Bornstein et al.’s (2023) study, however, was to show that the start-point setting process is itself one of evidence accumulation. To do this, they computed a metric termed the *reinstatement index*, the integrated evidence of stimulus reinstatement over the cue period, measured at each timepoint as the correlation between multi-voxel patterns measured when subjects directly perceived each target image with those observed when subjects engaged in putative cue-based retrieval of that target. Reinstatement indices were computed for neural activity in two regions of interest only—the fusiform face area (FFA) and the parahippocampal place area (PPA)—whose activity is known to be specialized for discriminating similar faces and scenes, respectively. Importantly, the authors theorized that the reinstatement index should be *inversely* related to associative memory strength: weaker associations require more sampling from memory, whereas stronger associations can be retrieved after a smaller number of memory samples (Bornstein et al., 2023; Bornstein and Daw, 2013). Consistent with this hypothesis, Bornstein et al. (2023) found that the reinstatement index was lower on trials with a stronger predictive probability and increased as a function of time between onset of cue and response time. Interestingly, the authors also found that the reinstatement index was greater on trials with a long cue-stimulus ISI *only* when a cue signaled weak upcoming sensory evidence. The authors offered no formal explanation for this phenomenon, which we term *uncertainty-adaptive anticipatory memory sampling*.

#### 5.4.2 Computational modeling

The authors did, however, test variants of a diffusion model to investigate their hypothesis that expectation-setting proceeds via evidence accumulation. Their core model (multi-stage DDM; MSDDM) was comprised of *two* diffusion processes with independent drift rates, one corresponding to an expectation-setting phase and the other corresponding to the sensory accumulation phase. Importantly, the terminal value of the decision variable in the first process sets the starting point for the sensory accumulation process on a trial-by-trial basis, effectively positing a two-stage process whereby memory dynamically biases sensory evidence accumulation on a trial-by-trial basis (Bornstein et al., 2023; Srivastava et al., 2017; (Figure 6A, left)). The model assumes one common threshold for the whole process, and was tested against two other models that selectively disabled key features of the MSDDM: a standard DDM with a constant drift rate across trials (1DDM) and two different DDMs that were fit independently to the expectation-setting and sensory-accumulation phases of each trial (2DDM), critically *without* information passing from the first stage to the next. These comparison models thus allowed for testing the necessity of (1) a time-varying drift rate and (2) a direct link between expectation-setting and sensory-accumulation processes, respectively.

**Figure 6:**
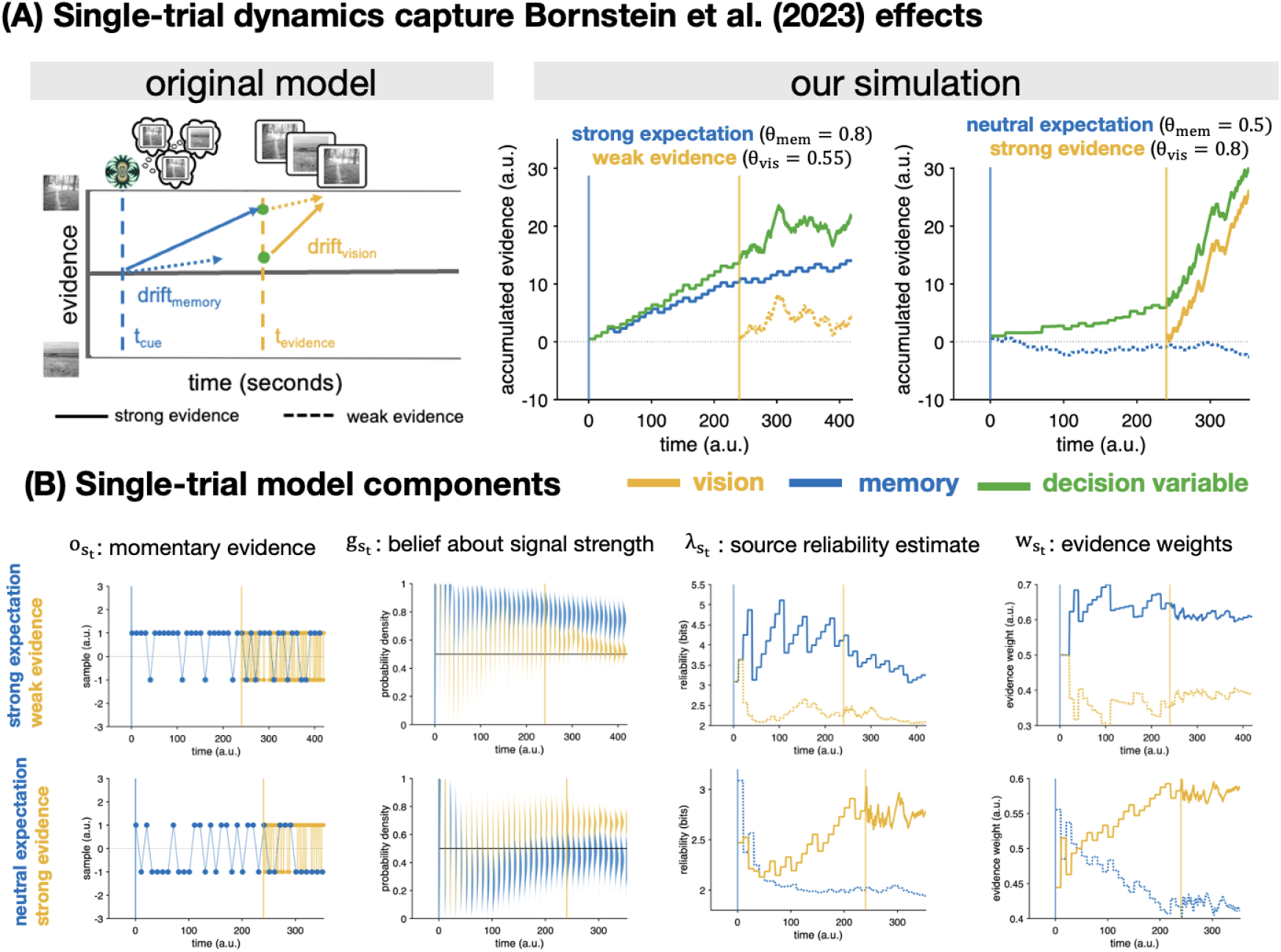
Dynamic reliability-weighted integration captures time-varying effects from Bornstein et al. (2023). In all figures, memory evidence (expectations; Π) is plotted in blue, vision evidence is plotted in yellow, and the decision variable is plotted in green. Vertical lines correspond to timepoints when memory (blue) and sensory (yellow) evidence come “online” for the observer. Memory samples in both simulations were generated at a rate of 6Hz (*γ* = 10). (A) Our model captures key dynamics produced by Bornstein et al.’s (2023) multistage DDM (left). First, we show that expectations set the starting point for sensory evidence accumulation in a trial-by-trial manner (right, both figures). Then, we show that anticipatory sampling from a strong expectation expedites the decision process once weak sensory evidence becomes available (left panel under “our simulation”). Additionally, we show that sampling from memory continues throughout the anticipation period when an expectation carries no information about the likely choice outcome, such that choice dynamics are dominated by the strong sensory evidence once it comes online (right panel under “our simulation”). (B) Single-trial model components that generated the left and right panels for the “our simulation” portion of A. Components for the left panel—strong expectation with weak evidence—are plotted in the top row, and components for the right panel—weak expectation with strong evidence—are plotted in the bottom row.

To link the reinstatement index with predictions of the MSDDM, Bornstein et al. (2023) quantified the relationship between the reinstatement index and response times for choices made after the onset of sensory information. First, the authors found that the reinstatement index predicted faster response times even when controlling for cue strength or evidence coherence, indicating that this metric captures meaningful trial-level variability in choice behavior. Next, Bornstein et al. (2023) found that this speeding was unique to trials where the cue validly predicted the choice outcome, consistent with a hypothesis that the reinstatement index corresponds to a start-point setting process. Finally, the authors found that the effects of reinstatement index were strongest for valid trials with weak sensory evidence. Taken together, these findings strongly support the idea that humans use their expectations in an uncertainty-adaptive manner: drawing more strongly on internal evidence when sensory evidence is insufficient to make the decision, but not bothering to do the extra effort of memory retrieval when decisions are likely to be easy to resolve with sensory evidence alone.

#### 5.4.3 Simulation methods

We used both trial-level and steady-state simulations to demonstrate our model’s ability to capture the uncertainty-adaptive anticipatory sampling reported by Bornstein et al. (2023). Sensory evidence was generated at 60 Hz, and memory evidence was generated at frequencies ranging from 2-12 Hz (*γ* ∈ {30, 20, 15, 12, 10, 8, 7, 6, 5} and with evidence encoding noise *ɛ* ∈ {0.01, 0.06, 0.11, 0.15, 0.2}. We fixed the threshold to an arbitrarily large value of *z* = 250 because, in a fixed-period task, the dynamics we aimed to capture are largely independent of threshold. The authors tested expectation levels Π*_cue_* = {0.5, 0.6, 0.7, 0.8} and sensory evidence levels individually calibrated to two separate strengths corresponding to low (65%) and high (85%) decisional accuracy in the absence of any expectation (Bornstein et al., 2023). We focus on capturing the two dynamics that comprise the uncertainty-adaptive anticipatory sampling finding: a strong expectation predicting a low accuracy (i.e., weak evidence) trial, and a neutral expectation predicting a high accuracy (i.e., stronger evidence) trial. The corresponding steady-state timeseries were generated using the same process described in Section 4.2. As in Bornstein et al. (2023), we fixed the duration of sensory evidence presentation to 3000ms and ran separate simulations for each of the short, medium, and long anticipation durations (4000, 6000, and 8000ms) used in the study. Main text findings report simulations from the shortest cue-stimulus interval (4000ms); steady-state dynamics for the longer two intervals can be found in Supplemental Figure **??**.

Additionally, to capture the joint nature of cues in Bornstein et al. (2023), we added *anticipated* coherence to the relative reliability computations occurring during the cue-stimulus interval. Rather than set sensory evidence weights to zero, we assume that each new sample of evidence from memory gives observers information how reliable the upcoming sensory signal will be. Specifically, for each timestep where a memory sample is drawn before the onset of sensory evidence, simulated agents receive one noisy sample from a Bernoulli distribution where *θ_vis_* is defined by the sensory signal reliability on that trial. Importantly, this sample is used only to update the belief about upcoming signal reliability *g*(*t*)*_viz_*, the time-varying reliability of each evidence source *λ*(*t*)*_s_*, and the time-varying weights on evidence *w*(*t*)*_s_* defined by the relative values of *λ_s_* at each point in time; it does not enter into either the sensory accumulator or the decision variable.

We do, however, assume that the decision variable is continuously updated with each new piece of information from memory during the period of time between the onset of the cue and the onset of sensory evidence. Specifically, we update the decision variable as follows:

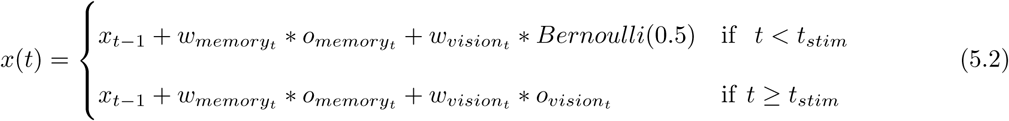

where *t_stim_* denotes the timepoint at which sensory evidence becomes available.

#### 5.4.4 Simulation results

Figure 6A shows that single-trial dynamics of our model (right) capture key properties of Bornstein et al.’s (2023) MSDDM (left). First, both panels in Figure 6A (right) show that our model creates a process where the memory accumulator (blue lines) sets the starting point for the vision accumulator (yellow lines) on a trial-by-trial basis. When a strong expectation about choice outcome *also* predicts weak upcoming sensory evidence (first panel, right side of Figure 6A), the decision variable (green line) is increasingly biased toward the choice predicted by the expectation as a function of elapsed time between cue onset (vertical blue line) and sensory evidence onset (yellow vertical line). However, when an expectation carries no information about likely choice outcome, but signals strong upcoming sensory evidence (second panel, right side of Figure 6A), the decision variable is much less biased by memory during the interval between cue and sensory evidence. Figure 6B displays the corresponding model components for these single-trial dynamics. When memory evidence is consistently more reliable than sensory evidence (top row), the model continuously places more weight on memory than vision throughout the trial (last panel, top row). However, when memory is uninformative, the model switches to placing more weight on sensory evidence *during* the anticipation period (last panel, bottom row). If we consider weights on each evidence source as corresponding to observers’ incentive to sample information from that source (as we did in Section 5.2), then these dynamics can be interpreted as reproducing Bornstein et al.’s (2023) observation uncertainty-adaptive anticipatory sampling effect on the single-trial level.

Figure 7 shows that uncertainty-adaptive anticipatory sampling dynamics are also present in steady-state model simulations. The effect is captured by difference between the solid and dashed lines in the right panel of Figure 7A: weights on informative expectations increase with elapsed time between cue and stimulus onset, and the magnitude of these weights is greater when sensory evidence is anticipated to be highly noisy (dashed blue line vs. solid blue line). The steady-state source reliability estimates displayed in Figure 7B further elucidate the time-evolving quantities driving these dynamics. For timepoints before the onset of sensory information (*x* ≈ 225), reliability estimates are formed on the basis of an *expected* value about sensory evidence strength that is estimated at the frequency of the memory sampling process. These slower evidence generation dynamics lead to inflated source reliability estimates, as discussed in Section 4.2. Once sensory evidence actually comes online and presents the observer with far more samples per unit time, the reliability estimate *λ_vis_* quickly reaches its asymptotic value, which is less than the original estimates based on memory samples. Memory reliability, however, continues to be estimated at the same rate throughout the decision and thus remains steadily elevated above the reliability of vision when the difference in *θ_s_* is sufficiently large.

**Figure 7:**
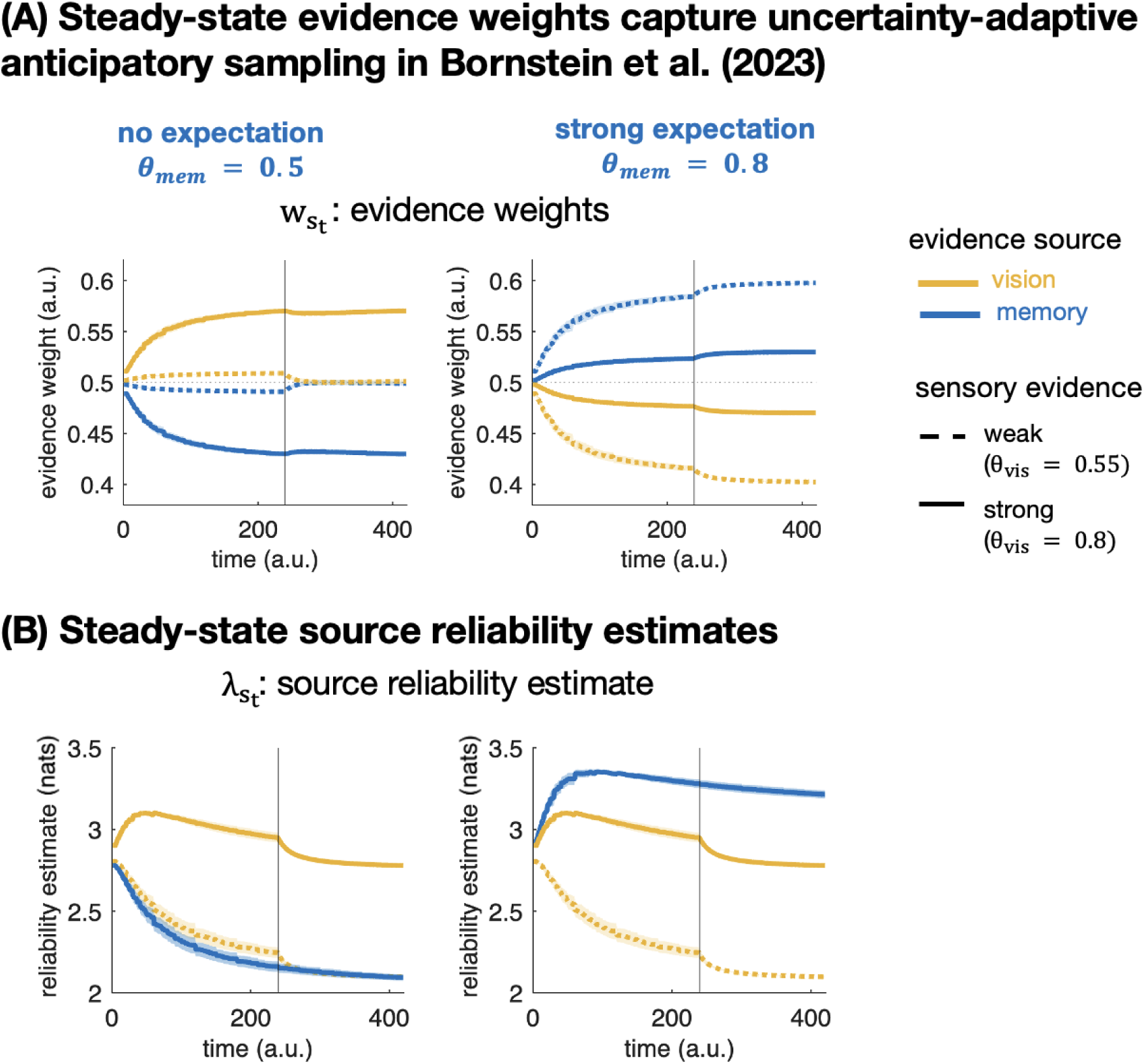
Steady-state dynamics of evidence weights and source reliability estimates driving Bornstein et al.’s (2023) observation of uncertainty-adaptive anticipatory memory sampling. Quantities corresponding to memory are plotted in blue and quantities corresponding to vision are plotted in yellow. Solid lines correspond to simulations with a weak sensory signal (*θ_vis_* = 0.55), dashed lines correspond to simulations with a strong sensory signal (*θ_vis_* = 0.8). The vertical line indicates the onset of sensory evidence after a 4000ms anticipation period; dynamics for longer anticipation durations are plotted in Figure **??**. (A) Weights on each evidence source throughout a single decision. (B) Source reliability estimates throughout a single decision.

The steady-state evidence weights for the comparison case of an uninformative expectation paired with strong sensory evidence (left panel of 7A) are interestingly non-monotonic. When a cue makes no prediction about the likely choice outcome, but signals that upcoming sensory evidence will be strong (solid lines), increasingly more weight is placed on vision relative to memory during the cue-stimulus interval. However, once the sensory evidence becomes available (vertical line), the model slightly and briefly *downweights* vision immediately after evidence onset, before returning to the same dynamics of increasing the weight on vision relative to memory. This transient downweighting is also observed—albeit to a lesser degree—for longer cue-stimulus interval durations (Figure **??**). This downweighting is again explained by examining the reliability estimates in left panel of Figure 7B. Because weighs on vision are initially generated based on an *expectation* about *θ_vis_*, and because that expectation generates information at the theta oscillatory frequency (2-12Hz), vision reliability is *overestimated* relative to its asymptotic value. In this case, however, the weights on sensory evidence (top panel), *increase* again after an initial “dip” during which *λ_vis_* reaches asymptote.

Taken together, both the single-trial and steady-state simulations in this section demonstrate that our model can capture the core time-varying effect observed by Bornstein et al. (2023): uncertainty-adaptive anticipatory sampling of learned expectations on a trial-by-trial basis. Additionally, we showed that our parallel process generates the same trial-level start-point setting specified by Bornstein et al.’s 2023 MSDDM, suggesting that our model might be a more general case of the two-stage process they identify.

### 5.5 Quadrant IV: trial-level explicit expectations (Hachisuka, Shor, Liu et al., 2026)

A recent study by Hachisuka, Shor, Liu et al. (2026)measured human behavior and neural activity in experimental settings that correspond to the fourth and final quadrant in Figure 1: an expectation-heterogeneous environment where observers have uncertainty about *which* expectation to use for each consecutive choice, but little to no uncertainty about *what* each expectation predicts.

#### 5.5.1 Study design & key findings

Hachisuka, Shor, Liu et al. (2026)combined high resolution fMRI, intracranial recordings, and deep neural network modeling to investigate the neural and computational mechanisms of one-shot perceptual learning in humans. To do this, they used “Mooney images”: photographic stimuli that initially appear ambiguous or abstract, but are actually just degraded versions of images that are instantly recognizable when presented without degradation (examples are shown in Figure 8A). Crucially, the apparent ambiguity of Mooney images disappears after just one exposure to the full-resolution image, such that observers cannot “unsee” the true contents of degraded image after being exposed to its full-resolution counterpart. This property of Mooney images makes them well-suited for studying the neurocomputational mechanisms involved in one-shot perceptual learning.

**Figure 8:**
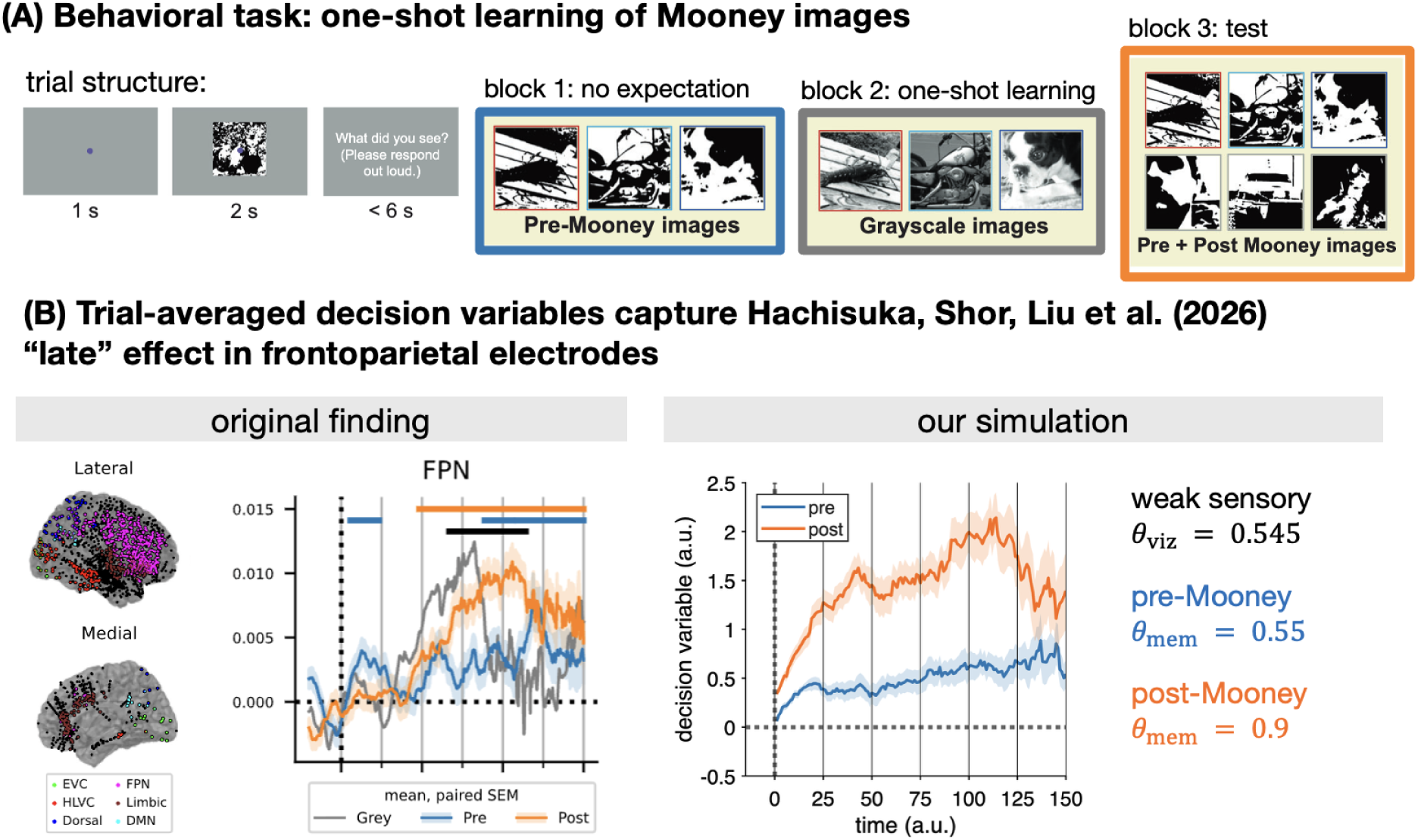
Dynamic reliability-weighted integration captures Hachisuka, Shor, Liu et al. (2026)“late” effect of expectations. (A) Experimental design in Hachisuka, Shor, Liu et al. (2026). On each trial, human subjects were presented with an image for 2 seconds and asked to verbally report the content of the image. In the first block, participants viewed “degraded” images whose contents are difficult to discern in the absence of prior knowledge. Then, in a second block, participants were presented with denoised grayscale versions of the same images, facilitating the one-shot learning characteristic of Mooney images. Finally, in the third block, participants were presented with degraded images again. Crucially, 50% of the images had just been presented in full resolution during the learning block. This manipulation allows for discriminating neural timecourses as a function of preand post-learning. Image reproduced from Hachisuka, Shor, Liu et al. (2026)with colored borders added for clarity. (B) Original finding: location of implanted electrodes across all patients (left), average activity from frontoparietal (FPN) electrodes (right). Activity post-learning (orange) ramps up more quickly than activity pre-learning (blue), generating a “late” effect of one-shot learned expectations about ambiguous sensory input. Our simulation: decision variables averaged across trials reproduce the pre- and post-learning dynamics measured by FPN electrodes. Shaded areas reflect standard error across *n* = 20 simulated subjects.

Hachisuka, Shor, Liu et al. (2026)developed a task where neural activity is recorded upon initially viewing a Mooney image (“pre-Mooney”), then viewing the full-resolution corresponding image in grayscale (“gray”), and then finally viewing the Mooney images again after having seen their full-resolution counterparts (“post-Mooney”; Figure 8A, right). On our framework, this corresponds to an initial “no expectation” block (blue border, Figure 8A), followed by a “one-shot learning” block (gray border, Figure 8A), and then finally a “test” block (orange border, Figure 8A). In each block, trials began with a fixation cross presented for 1000-2000ms followed by presentation of a Mooney or grayscale image for 2000ms (Figure 8A, left). Patients with intracranial electrodes were prompted to answer the question “Can you name the object hidden in the image?” with Yes/No keyboard presses made using different hands, whereas non-patients were prompted to verbally respond to the question “What did you see?”. Crucially, the test block following grayscale image presentation contained Mooney images observed in the first block (now *post*-Mooney) as well as brand new “pre”-Mooney images that had not been previously presented to the observer. Thus, the “post” images in the test block are those for which observers have an unambiguous expectation, whereas there is no such expectation for the “pre” images.

The authors report cross-modal evidence suggesting that one-shot perceptual expectations are encoded in higher-level visual cortex (Hachisuka, Shor, Liu et al., 2026). The main finding of interest for the present analysis, however, comes from intracranial recordings from the frontoparietal network (FPN), which encompasses several key regions implicated in perceptual decision-making (Hanks and Summerfield, 2017; Polanía et al., 2014). As shown in the left panel Figure 8B (“original finding”), FPN neurons exhibited accumulation-like responses to the Mooney images, with responses to post-Mooney images (orange line) increasing nonlinearly over time relative to pre-Mooney images (blue line). The authors suggest that this difference in slope later in the trial “may be related to recognition-triggered decision-related activity” (Hachisuka, Shor, Liu et al., 2026, p.7), which is precisely what our model predicts ought to occur. To test the ability of our model to capture these dynamics, we conducted the simulations described below.

#### 5.5.2 Simulation methods

The original timeseries data were generated from approximately 500 FPN electrodes implanted across *n* = 19 patients. Each patient completed at least 30 trials of the pre/post Mooney task, and timeseries were averaged across trials in each condition for each electrode in each subject (Hachisuka, Shor, Liu et al., 2026). Because the form of our model is designed to capture individual-level processes, we treated each run of the simulation as the average of all electrodes implanted in a single patient. Further, we assumed that each subject contributed data using some unique contribution of relative memory “sampling rate” *γ* and threshold *z*. For all subjects, we defined the strength of sensory and memory evidence as follows: *θ_vision_* = 0.545, *θ_pre_* = 0.55, and *θ_post_* = 0.9. We chose values slightly greater than the minimum of 0.5 for sensory evidence and pre-Mooney expectation to capture the effects of low-level visual signals and repetition-based familiarity, respectively (Hachisuka, Shor, Liu et al., 2026). As in the original study, sensory evidence was generated at a rate of 75Hz and presented for 2000 simulated ms (*n* = 150 maximum timesteps per trial).

Given the rate of sensory evidence presentation, we generated data using *γ* = {6, 7, 8, 9, 10, 12, 15, 18, 21, 30} such that memory samples were generated at rates spanning 2.5-12.5Hz across all subjects. The decision process was terminated using threshold values *z* = {4, 6}. As in Section 5.2, we manipulated evidence encoding noise *ɛ* on the whole-sample level, running different simulations for *ɛ* ∈ {0.01, 0.06, 0.11, 0.15, 0.2}. As with all other findings, we report results from the simulation with *ɛ* = 0.11; timeseries averaged across all *ɛ* values are displayed in Figure **??**. For each cue type (pre-vs. post-Mooney), we simulated 30 trials using each unique combination of *γ* and *z*, which created a sample size of *n* = 20 simulated subjects. We averaged the decision variable of our model across runs of the simulation within each subject, and then display the group averages in Figure **??**B, which also contains shaded regions depicting the standard error across simulated subjects.

#### 5.5.3 Simulation results

We focused our simulation only on the pre- and post-Mooney timeseries (blue and orange lines in Figure 8B), since these recordings are the most relevant to our theory. The pre-Mooney timeseries correspond to accumulation of weak sensory evidence with only a weak expectation about its content, whereas the post-Mooney timeseries correspond to accumulation of the same weak sensory evidence but with a strong expectation about its content. Importantly, observers are not told *which* grayscale image to use as their expectation for any given trial in the test block; they simply observe the Mooney image and then make a report.

Our sampling-based theory of expectation retrieval, setting, and integration predicts that time-varying effects of memory retrieval on choice dynamics ought to be observed in precisely this kind of experimental setting. Figure 8B quantitatively validates that prediction by showing that our dynamic reliability-weighted process can reproduce similar dynamics as those observed by Hachisuka, Shor, Liu et al. (2026)in neural recordings. When observers have learned a strong expectation for a particular sensory input, but are not instructed ahead of time to deploy that expectation for an upcoming choice, then their choices become increasingly biased by that expectation as a function of elapsed decision time (orange lines in Figure 8B). However, when an observer does *not* have the relevant expectation for a particular sensory input, the decision variable increases at a lower and more constant rate (blue lines in Figure 8B). Figure **??** further demonstrates that these effects are present in steady-state dynamics of our model as well.

Hachisuka, Shor, Liu et al. (2026)speculated that these time-varying dynamics might be driven by recognition-related neural activity. Although we do not explicitly include recognition-driven retrieval dynamics in our model, our ability to reproduce the “late” effect observed by Hachisuka, Shor, Liu et al. (2026)lends support to the authors’ original interpretation of memory-related dynamics. Further, we offer a formal account for *why* these dynamics occur *when* they do: expectation-setting takes time, and uncertainty about *which* expectation to use across decisions will increase the amount of time it takes to set & integrate the relevant expectation into the decision process. These recent findings from Hachisuka, Shor, Liu et al. (2026)nicely complement the late effect of expectations first reported by Hanks et al. (2011). Whereas the late effect in Hanks et al. (2011) was driven by heterogeneity in the sensory evidence strength across trials, the late effect in Hachisuka, Shor, Liu et al. (2026)is driven by heterogeneity in the *expectation* across trials. While it is possible that some Mooney images are more easily discernible than others in the pre-learning stage, this incidental variability in sensory evidence strength is unlikely to drive robust effects across observers. Taken together, the Hanks et al. (2011) and Hachisuka, Shor, Liu et al. (2026)findings lend converging support for the core theoretical argument of this paper: dynamic effects of expectations are driven by the time-varying *relative* uncertainty of memory and sensory evidence.

### 5.6 Interim summary

The previous two subsections (5.4 and 5.5) show that our model can reproduce dynamic effects of expectations observed in *expectation-heterogeneous* environments, where observers face deployment-related uncertainty about which expectation will be used on each trial (left side of the quadrants in Figure 1). First, we reproduced the uncertainty-adaptive anticipatory sampling finding reported by Bornstein et al. (2023), where observers increase their sampling of memory over elapsed time before evidence onset *only* when a predictive cue signals weak upcoming sensory evidence (Figure 6A, right; Figure 7A, right). Then, we generated timeseries that reproduce the difference in pre- and post-learning activity observed in frontoparietal electrodes while humans discriminated Mooney images (Hachisuka, Shor, Liu et al., 2026), lending formal support to the original authors’ interpretation of memory-related processing.

Combined with earlier findings that our model captures captures dynamic effects in expectation-*homogeneous* environments (Sections 5.1-5.2), this section has shown that our principled model can reproduce time-varying effects of expectations on perceptual decisions observed across different experiments, species, neuroimaging modalities, and research groups. Each of these findings were initially explained using distinct cognitive and/or computational processes: learning time-dependent accuracy functions (Hanks et al., 2011), conflict monitoring (Rungratsameetaweemana, Itthipuripat et al., 2018), two-stage evidence accumulation (Bornstein et al., 2023), and memory recognition (Hachisuka, Shor, Liu et al., 2026). What we have shown, however, is that these findings can be unified through our principled theory of uncertainty-driven memory sampling and integration (Figures 1 and 2).

Importantly, our results do not necessarily imply that previous explanations are incorrect. Indeed, most of them already invoke different theoretical dimensions of our model: uncertainty-driven dynamic weighting on expectations (Hanks et al., 2011), expectation-setting via evidence accumulation (Bornstein et al., 2023), and memory retrieval/reinstatement as a process that contributes evidence to the perceptual decision variable (Hachisuka, Shor, Liu et al., 2026). Our model can thus be interpreted as *synthesizing* existing theories about time-varying effects through a principled computational framework, rather than invalidating the previous findings or explanations.

### 5.7 Uncertainty about expectations induces sequential effects on RTs

In addition to unifying previously-disparate observations into a common principled process, the framework of uncertainty-driven memory sampling generates a testable empirical prediction: if the process of *setting* expectations itself involves evidence accumulation over time, then *switching* expectations across consecutive decisions ought to *slow* reaction times relative to when expectations remain unchanged across decisions. Further, this effect should be observed independently of other behavioral and cognitive factors known to modulate reaction time. We tested this prediction using linear regressions on publicly-available data from two studies that measured human behavior in expectation-heterogeneous environments, but differed in whether observers used learned cues through instruction (Diaz, Pisauro et al., 2024)or experience (Bornstein et al. (2023)); details about each study can be found in Section **??**.

Regressions were fit using the lm() function in R 4.4.2. Estimated marginal means, 95% confidence intervals, and contrast coefficients were computed using the emmeans package. All reaction times were log-transformed and z-scored within subject before entered into the regression, and thus we did not include any random effects for subjects. All models contained fixed effects of predictors of interest (previous cue and previous target), as well as for factors known to affect reaction time (cue validity, cue level, choice accuracy, sensory evidence strength, accuracy of choice on immediately preceding trial, and total amount of time on task). Sensory evidence strength and time on task (defined as raw trial number) were modeled as continuous regressors, whereas all other variables were modeled categorically so as not to enforce a linearly-increasing effect of cue probability. Data and code to reproduce these results can be obtained from Github.

Figure 9 shows that both instructed and learned expectations induce sequential effects on reaction times. For both the Diaz, Pisauro et al. (2024)and Bornstein et al. (2023) datasets, RTs were significantly slower when cues switched across two consecutive trials relative to when they were repeated (*β_instructed_* = 0.029, *t*_16687_ = 3.858, *p <.*001; *β_learned_* = 0.055, *t*_4678_ = 3.172, *p* =.002). Crucially, we did not observe any significant effects of switching or repeating choice targets across trials (*β_instructed_* = 0.003, *t*_16687_ = 0.516, *p* = 0.606; *β_learned_* = −0.007, *t*_4678_ = −0.363, *p* =.717) in either dataset, suggesting that the observed effects are driven solely by switching expectations—but not sensory choice outcomes—across trials. Full summaries of fitted coefficients can be found in Section **??**.

**Figure 9:**
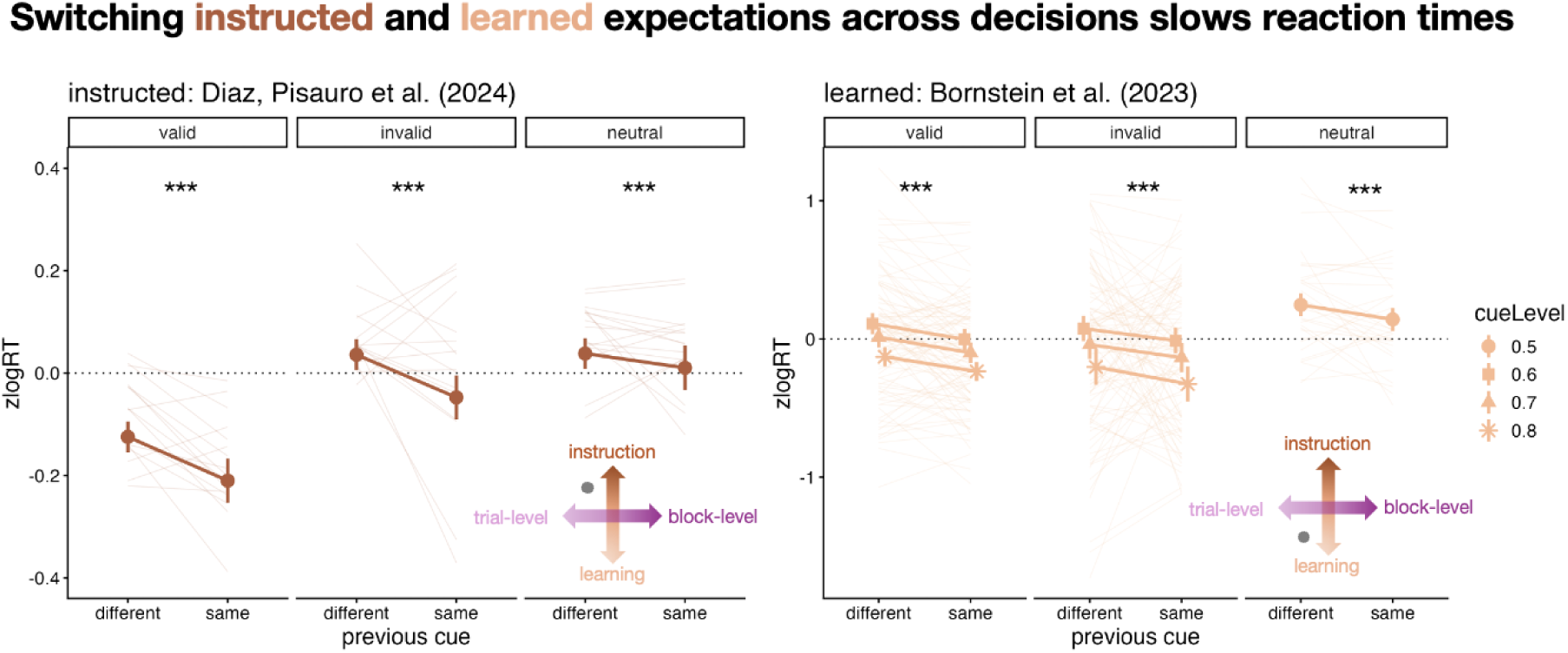
Expectation-heterogeneity induces sequential effects on RTs, regardless of whether expectations are explicitly instructed (left) or learned through experience (right). Across all cue levels, switching cues across trials significantly slows responses, whereas repeating cues across trials significantly speeds responses. In all panels, individual lines reflect subject means, points are group-level marginal means estimated using fitted regression coefficients, errorbars reflect 95% confidence intervals around that estimate.

Taken together, these findings lend empirical support to the core claim of this section: uncertainty about expectations ought to induce effects on the dynamics of perceptual choice. We focused our analyses on behavior measured in expectation-heterogeneous environments, where observers have uncertainty about *which* expectation they will need to use for each decision. Our uncertainty-driven memory sampling framework predicts that behavior should exhibit “switch costs” related to the time it takes to adjust expectations across trials with different cues, which is precisely what we observed in both the Diaz, Pisauro et al. (2024)and Bornstein et al. (2023) datasets. Further, these effects were observed independently of canonical experience-related effects on RT, providing evidence for effects of uncertainty at both the trial- and task-level. These findings demonstrate the theoretical utility of considering how uncertainty about expectations modulates the dynamics of perceptual choice.

## 6 Discussion

We have shown that uncertainty-driven memory sampling can explain time-varying effects of expectations on perceptual decision-making. A core theme throughout the article has been interrogating the assumptions behind extant models specifying the optimal procedure for integrating beliefs about prior probability (or expectations) into evidence accumulation processes. In doing so, we identified *uncertainty about expectations* as a critical feature that has been omitted from existing normative models. We then used the framework of sequential sampling to develop a verbal theory and formal model articulating how uncertainty about expectations ought to impact evidence accumulation. Regression analyses on existing behavioral data provided empirical support for a novel prediction generated by our verbal theory, and simulation results demonstrated that our model can reproduce time-varying effects of expectations observed under various experimental conditions and neuroimaging modalities.

This section further articulates the contributions that our framework and model make to broader literature on expectations in perceptual decision-making. To do this, we first discuss how our model is related to—and distinct from—neighboring models in the literature. Then, we discuss insights our model offers for neural investigations into evidence accumulation. Next, we discuss the simplifying assumptions upon which our model rests, articulating both how those assumptions constrain the scope of phenomena the model can capture and how they might be relaxed to capture more complex phenomena of interest. Finally, we conclude with novel predictions that our model makes about choice behavior and neural dynamics, further demonstrating its utility for advancing research on memory-guided visual behavior.

### 6.1 Related models

The sequential sampling family of models is broad and structurally heterogeneous, with each model specifying a slightly different process whereby evidence is accumulated to commit to a choice. Further, different models within this family have been developed to support different approaches to scientifically studying decision making. We have previously shown how considering the difference in reasoning goals motivating model specification and application can resolve apparent tensions in model comparison and selection, especially concerning the inclusion of components that vary as a function of time within the trial (Khoudary et al., 2025b).

In this work, we use sequential sampling to specify a formal theory about a data-generating process whereby two sources of dynamic, probabilistically-evolving evidence are combined based on first principles from statistics and psychology (i.e., reliability-weighted cue combination). We structure our discussion of related models along this axis of use cases, beginning first by addressing other simulation-based findings of time-varying diffusion models. Then, we discuss “multi-stage” diffusion models that are often used to approximate continuously-changing processes when models are applied to data. Finally, we discuss “psychometric” diffusion models, a term we use to refer to models designed specifically for purposes of latent variable estimation.

#### 6.1.1 Dynamic diffusion models

The first class of models we survey are those where the drift rate and/or decision threshold change continuously as a function of time (Deneve, 2012; Huang et al., 2012; Malhotra et al., 2017). One of the most similar previously-specified models was proposed by Deneve (2012), who investigated the question of how the dynamic bias signal identified by Hanks et al. (2011) might be implemented via Bayesian neural inference. In addition to inferring the probability of the correct choice conditional on all observed evidence thus far, Deneve’s (2012) model also infers the sensory evidence strength on a trial-by-trial basis. The model then uses this time-varying estimate to scale the decision thresholds and weighting of evidence within the course of a single decision. This results in a process where both sensory weights and decision thresholds *increase* with time on trials with highly certain sensory evidence, but *decrease* with time on trials with highly *uncertain* sensory evidence (Deneve, 2012, Figure 3). Deneve (2012) further shows that this process also generates time-varying weights on expectations, such that they become more strongly weighted in the decision process as a function of elapsed time on a single choice. Finally, Deneve (2012) shows how these computations can be implemented using both spike trains of single motion-selective neurons as well as populations of neurons that share motion-selective properties, offering proof-of-principle for the feasibility of (1) jointly inferring uncertainty and motion direction and (2) using these quantities to dynamically modulate decision boundaries and drift rates (i.e., weighting of evidence at each moment in time).

Complementary analyses by Huang et al. (2012) and Malhotra et al. (2017) used partially observable Markov decision processes (POMDPs) to further demonstrate that the dynamic bias signal reported by Hanks et al. (2011) can be generated via time-varying decision boundaries. As discussed in Section 3, POMDPs permit framing the evidence accumulation process as one of action selection: at each moment in time, an observer chooses whether to continue sampling or commit to one of the two choice outcomes based on the evidence they have accumulated thus far (Equation 3.1). The policy of a POMDP defines which action the observer takes at each timestep, and optimal policies are those that maximize expected future reward (Rao, 2010; Huang et al., 2012; Malhotra et al., 2017). Huang et al. (2012) showed that the optimal policy for a POMDP in an expectation-homogeneous, evidence-heterogeneous environment corresponds to decision boundaries that (1) decrease exponentially over time and (2) are shifted in proportion to the prior probability Π, effectively implementing a starting point offset. A more recent study from Malhotra et al. (2018) built upon these findings to show that the *rate* of bound collapse depends jointly on the prior probability and the mixture of sensory evidence strength in the decision environment. The authors first showed that decision boundaries only exhibit exponential collapse in environments where one of the signal strengths is sufficiently low; otherwise, reward rate is optimized by using boundaries that are constant over time (Malhotra et al., 2018). Then, they showed that adding a strong prior probability *linearizes* the collapse of boundaries over time (Figure 9 in Malhotra et al., 2018), such that for sufficiently high prior probability (in this case, Π = 0.7) the optimal boundaries remain parallel but decrease monotonically over time. Accordingly, increasingly more evidence is required to overcome the prior as deliberation time increases—precisely the dynamic effect observed by Hanks et al. (2011) and reproduced by our model.

Taken together, the simulation-based findings of dynamic diffusion models strongly suggests that a time-varying modification to the decision process is needed when observers have uncertainty about sensory evidence (Drugowitsch et al., 2012; Deneve, 2012; Huang et al., 2012; Malhotra et al., 2018). Although our model does not explicitly incorporate time-varying decision boundaries, we have shown that it reproduces both the original finding these models aimed to explain (Hanks et al., 2011) in addition to explaining several other time-varying effects of expectations on perceptual decisions (Section 5). Critically, our model suggests that several of the dynamics predicted by expectation-induced effects on decision boundaries could also be generated by a dynamic reliability-weighted process integrating samples from parallel streams of memory and sensory evidence. Because time-varying drift rates can often mimic time-varying thresholds, carefully designed experiments coupled with high-resolution neuroimaging will likely be required to adjudicate these formally similar—but conceptually different—theories of how expectations modify the dynamics of choice. Further, as Deneve (2012) suggests, it could be the case that expectations exert time-varying effects on *both* the drift rate and decision thresholds. One possibility is that expectations with low source-related uncertainty act primarily on decision thresholds, whereas expectations with high source-related uncertainty act primarily on the drift rate, owing to this latter parameter’s weaker dependence on observers’ explicit knowledge or decisional strategies.

#### 6.1.2 Multi-stage diffusion models

Multi-stage diffusion models correspond to a stochastic process where the drift rate, diffusion coefficient, and threshold change discretely—rather than continuously—over time (Srivastava et al., 2017). We focus specifically on the class of multi-stage *drift* models, which generate “switches” in the slope and direction of the decision variable at specific points in time (Ratcliff, 1980; Diederich and Busemeyer, 2006; Diederich and Oswald, 2014, 2016; Srivastava et al., 2017).

##### 6.1.2.1 Two-stage models of expectations

A study by Diederich and Busemeyer (2006) was among the first to apply this kind of model to the question of how asymmetries in response outcomes—defined as differences in expected payoffs—modulate choice dynamics. The authors found that a two-stage model where payoffs were initially processed with one drift rate, and then stimuli were processed with a different drift rate outperformed models that incorporated the payoffs as offsets on both the starting point and drift rate (Diederich and Busemeyer, 2006). This finding is a conceptual predecessor to the results of Bornstein et al. (2023), who used a model developed by Srivastava et al. (2017) to show that brain and behavioral data were best captured by a two-stage process whereby evidence accumulation from memory sets the starting point for sensory evidence accumulation on a trial-by-trial basis.

The right panel of Figure 6A shows that the decision variable of our model effectively implements a time-continuous version of the two-stage model used by Bornstein et al. (2023): anticipatory accumulation from memory biases the starting point of the sensory evidence accumulator jointly as a function of expectation strength and sampling time. Two key features distinguish our model from the multi-stage models developed by Srivastava et al. (2017) and Diederich and Busemeyer (2006). First, our model posits two independent accumulators for each evidence evidence source, and a decision variable that is a dynamic reliability-weighted combination of samples from each source. This leads to the second difference, which is that we posit an effect of memory that *continues on* once the sensory evidence has been presented. Rather than suggest that the decision process involves discrete “switches” based on sampling one source versus another, we propose that both sources of evidence are continuously sampled and integrated throughout the deliberation process. We discuss this assumption of our model, and possible extensions beyond it, in later parts of the Discussion.

##### 6.1.2.2 Opposing process theory

Although not expressed using the formalism of sequential sampling, Press et al.’s (2020) “opposing process theory” can be thought to correspond to a process with a multi-stage drift rate. Press et al. (2020) developed opposing process theory to resolve an apparent paradox between “Bayesian” and “cancellation” theories of how expectations modulate the gain of sensory information. Whereas Bayesian theories—prominent in sensory perception research—assert that expected information ought to have a higher gain, cancellation theories— prominent in action research—assert that *unexpected* information ought to have higher gain. Opposing process theory resolves this tension by positing a two-stage process whereby sensory evidence for expected outcomes initially is processed with a higher gain, but that surprising inputs (quantified using the Kullback-Leibler divergence of an observer’s posterior relative to their prior) generate increased gain later on during processing, potentially via phasic release of catecholamine (Press et al., 2020).

The findings we display in Figures 5A and 6B suggest that our model is capable of formalizing the dynamics predicted by opposing process theory using a time-continuous modeling framework. The right panel of Figure 5A shows that our model generates greater weights on unexpected, followed by *unexpectable*, sensory evidence as a function of elapsed time. The rightmost panel of Figure 6B (top) additionally shows that strong expectations generate early weights that prioritize predictions from memory relative to contents of perception. Indeed, the conjunction of these dynamics—early upweighting of expected information and later upweighting of unexpected information—was recently observed via EEG decoding of a task specifically designed to test opposing process theory (Rittershofer et al., 2026). Given that our model can reproduce both of these dynamics in independent cases, it follows that it should be able to capture their conjunction as well.

#### 6.1.3 Psychometric diffusion models

The final class of models we consider are those that we term *psychometric* models, deriving from their development for purposes of latent variable estimation across a variety of experimental settings (Khoudary et al., 2025b; Batchelder, 2010). The first model we discuss, the attentional DDM (aDDM), posits a time-varying offset onto the decision variable as a function of fixation location (Krajbich et al., 2010; Tavares et al., 2017; Yang and Krajbich, 2023). The second, the extended DDM (eDDM), posits trial-level variability in the drift rate and starting point terms of the original DDM (Ratcliff and McKoon, 2008; Ratcliff et al., 2016). Our work in this article motivates alternative applications and/or interpretations of each model, which we discuss in turn.

##### 6.1.3.1 The attentional DDM (aDDM)

An increasingly prominent time-varying drift rate model is the “attentional DDM” (aDDM), which formalizes the effects of eye movements on the dynamics of value-based choice (2010). The aDDM posits that fixations on a specific option induce transient biases on the drift rate, such that evidence is accumulated more quickly for an option when an observer is overtly attending to it with their gaze. More recently, a multi-attribute generalization has been developed which further permits estimating the weighting of specific attributes—as opposed to entire outcomes—on the drift rate (Yang and Krajbich, 2023). A core feature distinguishing our model from the aDDM is *how* the dynamic effects on drift are generated: the aDDM uses measurements of fixations to *infer* time-varying biases on the drift rate, whereas our model posits that drift rate biases are a function of the time-evolving evidence itself. Further, the aDDM posits that the magnitude of drift modulations is governed by the subjective value of the attended choice option, whereas we suggest that the relative evidence reliability governs how strongly drift rates are biased.

These differences can be understood as reflecting the different cognitive processes—operationalized via different experimental tasks—that the models are designed to explain. In contrast to perceptual decision tasks that present one dynamically-evolving stimulus on each trial, stimuli in value-based decision tasks are comprised of two static, full-color images. Further, perceptual decision tasks have an objectively correct answer (defined via experimental manipulations), whereas value-based tasks are designed to elicit stable differences in subjective estimations of stimulus value. An interesting direction for future work would be developing a single model that unifies both types of decision processes into a common framework, perhaps by considering the role of uncertainty in scaling dynamic effects both of memory and attention on deliberation. Further, given the close empirical link between eye movements and memory retrieval (Barker et al., 2025; Kragel et al., 2020; Hannula et al., 2010), as well as memory retrieval and dynamic construction of subjective value (Shadlen and Shohamy, 2016; Wang et al., 2021), it would be fruitful to consider the role of memory retrieval dynamics in guiding fixations during both value-based and perceptual choice. In particular, it would be interesting to consider how memory sampling may dynamically *guide* eye movements during sensory evidence accumulation, perhaps by directing gaze to areas that in space that are maximally informative for discriminating whether a trial is congruent or incongruent with what is predicted by memory. Another possible direction for future research is using the aDDM to identify moments in time when observers’ attention switches from external sensory information to internal information generated by memory (Verschooren et al., 2026; Kumle et al., 2025), a process that has been linked to differential recruitment of the hippocampus by subcortical and cortical brain regions, respectively (Poskanzer and Aly, 2023).

##### 6.1.3.2 The extended DDM (eDDM)

The extended DDM (eDDM; Ratcliff and Rouder, 1998; Ratcliff and Tuerlinckx, 2002; Ratcliff and McKoon, 2008) builds upon the original DDM (Equation 2.1) in two ways. First, it specifies an additional *non-decision time* term *T*_0_ that additively offsets the first-passage time of the Gaussian walk to account for delays in RT attributable to putatively non-decision related processing, such as sensory evidence encoding and motor command execution. Second, it defines the drift rate, starting point, and non-decision time terms as *probability distributions* rather than constant values, such that the decision process can exhibit structured variability across trials. Both of these modifications enhance behavioral goodness-of-fit across several tasks, but a principled theory about *why* trial-level variability in these parameters (and of their specific probabilistic form) is still largely lacking.

Anticipatory, simultaneous, and/or post-decisional memory sampling may offer some insight into this question. Specifically, trial-level variability in the starting point may reflect individual differences in how quickly memory information comes to mind and/or is weighted relevant to the upcoming decision. Drift rate variability could also be explained by periodic “bursts” of internal evidence generated by memory that direct observers’ attention away from their external environment, contribute internal evidence bearing on the externally-oriented decision, or some combination thereof (e.g., Nicholas et al., 2026). As such, it may be the case that the eDDM improves goodness-of-fit not because it better captures facts about the data-generating process, but rather that it more flexibly accommodates stable individual differences that are not accounted for by simpler models. Although hierarchical Bayesian approaches to parameter estimation handle concerns about individual differences in a statistically elegant and principled manner, they do not on their own provide answers to the deeper theoretical question of *why* individuals exhibit the differences that they do. Process models based on first principles—coupled with tasks designed to tease apart structured deviations from their predictions—are powerful complementary tools toward this end.

### 6.2 Relationship to neural data

Principled process models have been highly useful for advancing research into the neural bases of evidence accumulation, and in linking neural activity with observed behavior across species (Brody and Hanks, 2016; Gold and Shadlen, 2007; Forstmann et al., 2016; Hanks and Summerfield, 2017). Foundational research in non-human primates demonstrated signatures of evidence accumulation in single-cell and population-level activity (Newsome et al., 1989; Britten et al., 1996; Gold and Shadlen, 2002), and mapped different computational steps of the formal process onto distinct cortical regions comprising a decision-making circuit (Gold and Shadlen, 2007). More recent research using large-scale recordings in rodents has both challenged and supported the early image developed based on primate work, primarily by demonstrating that signatures of evidence accumulation can be observed across the entire brain instead of a specialized decision circuit (Liu et al., 2025; Findling et al., 2025; Bondy et al., 2025).

The framework, model, and findings we develop here contribute to this line of research by identifying the specific circumstances in which memory retrieval dynamics are most likely to modulate the dynamics of sensory evidence accumulation. Further, our model permits making specific predictions about what a candidate timeseries might look like under different assumptions about the relative reliability of both evidence sources, the observer’s uncertainty about that reliability, and the relative rate at which information is retrieved from memory. And because we aimed to keep our model as minimal as possible, future work can refine its predictions by incorporating neural observations about the timescales of activity in different brain regions (e.g., Shi et al., 2025), which is a core conceptual idea motivating our model.

Another exciting direction for future research is to investigate alternative retrieval mechanisms to complement and extend our findings based on steady theta oscillations. One promising candidate to this end are sharp-wave ripples: brief, high-frequency bursts of hippocampal activity strongly associated with memory retrieval (Kerrén et al., 2026). Because our model does not restrict the type of mnemonic information that can contribute to decisions, it makes the prediction that *any* neural measurement of memory retrieval ought to display dynamic reliability-weighted effects. In direct support of this prediction, a recent study by Frank et al. (2026) reported that hippocampal ripples *increased* in response to visual uncertainty, and that this increase was directly associated with a decrease in reaction time. This observation is directly predicted by the model and framework we develop here, and is conceptually analogous to the uncertainty-adaptive memory sampling findings we model in Section 5.4.

### 6.3 Model assumptions and extensions

A core motivation for our model is identifying a principled process for integrating expectations with sensory evidence under less-idealized assumptions. Taking a formal approach, however, requires that we still make use of simplifying assumptions in order to make the modeling tractable. This section discusses these assumptions and suggests directions that future work can take to address them.

#### 6.3.1 Independence of memory and sensory evidence accumulation

First, our model assumes that each source generates and accumulates evidence independently of the other. That is, the generating distribution for memory samples remains constant over the course of a single decision and does not change as a function of the content of sensory information. We justified this assumption on the basis of our goal to model simple pattern completion and/or associative retrieval processes, rather than a more complicated memory search process. We believe this assumption of a static generating distribution for memory—or the independence of memory and sensory evidence samples—is warranted for quick perceptual decisions on the order of a few seconds. However, it is likely that more prolonged perceptual deliberation might involve sequentially searching through relevant information in memory (Wang et al., 2021; Callaway et al., 2024). Future work can investigate this possibility by dynamically manipulating the content of sensory evidence and/or the relevant expectation within the course of a single trial (e.g., Kiani et al., 2013; Kilpatrick et al., 2019), as well as considering cases where observers have more richly structured expectations about the dynamic contents of sensory evidence (e.g., Chen and Bornstein, 2024; Kaper and Peters, 2026).

#### 6.3.2 Memory “sampling rate” *γ*

Next, to capture putative differences in the timescales of sensory and mnemonic evidence generation, our model assumes that memory evidence is generated at a fraction of the rate that sensory evidence becomes available to the observer, captured by the *γ* parameter. For simulation findings aimed at reproducing empirical time-varying effects (Figures 5 and 8), we incorporated a wide range of memory “sampling rates” for two reasons: first, to show that our findings are robust across a range of plausible ratios in sampling rate, and second, to capture putative heterogeneity in this ratio across individuals and species (2014, 2020). We did, however, restrict the set of possible ratios to the broadest range of empirically-observed theta-band oscillatory activity (2-12Hz; Seger et al., 2023). While it is possible that the type of memory sampling we invoke here occurs at other relative “sampling rates”, we chose to restrict this set of findings to the theta frequency because of robust neural evidence implicating theta-band activity in task-evoked memory retrieval, particularly in the context of memory-guided visual behavior (Ter Wal et al., 2021; Zhang et al., 2018; Kragel et al., 2020; Kerrén et al., 2018; Zubair et al., 2026; Senoussi et al., 2025). An interesting direction for future research is to consider how memory “sampling rate” might be *controlled* by observers in a task- or uncertainty-dependent manner, such that information from memory can be made available more quickly on a case-by-case basis.

#### 6.3.3 Evidence reliability estimation

Our model makes a handful of assumptions about the reliability estimation process. First, we assume that observers dynamically estimate the reliability of each evidence source on every single decision; essentially, that observers treat the evidence-generating distributions as nonstationary with unknown time variation. This is a simplifying assumption because the evidence-generating distributions are commonly stationary at the single-trial level, so it is possible that reliability estimates may carry over from trial-to-trial. Indeed, the secondary behavioral analyses conducted in Section 5.7 corroborate the idea that reliability estimates are “cached” across trials, as evidenced by the RT advantage for repeating learned cues relative to switching them (Figure 9). Modeling and measuring how observers learn to estimate the reliability of time-evolving evidence—perhaps via a process akin to Kalman filtering (Roweis and Ghahramani, 1998)—is a promising direction for future research.

Second, and relatedly, we assume that observers estimate the reliability of evidence *online* during the decision process. This assumption follows directly from our aim of considering how expectation-heterogeneity— or uncertainty about *which* expectation to use on each trial—ought to impact the dynamics of perceptual choice: when the environment dictates which expectation to retrieve on a trial-by-trial basis, any estimation of mnemonic evidence reliability must be done during the retrieval process itself. It would be interesting, however, to consider cases where observers themselves determine which information to retrieve from memory, either in anticipation of or simultaneously with perceiving ambiguous sensory evidence. Our model predicts that observers ought to retrieve information that maximally reduces uncertainty in their deliberation process, but leaves open the question of *how* observers determine which information is useful toward this end. Tasks that utilize multidimensional expectations, along with sensory evidence that evolves along longer timescales, can help shed light on these questions (e.g., Finn et al., 2018; Nicholas and Mattar, 2026; Mazor et al., 2026). Further, it would be interesting to investigate whether the same dynamic effects emerge when evidence reliability is used to *guide* memory retrieval, rather than being estimated online as a function of it.

Finally, our model assumes that there is no noise or error in observers’ reliability estimation process. This assumption follows directly from Equation 4.3, which states that the Beta-distributed belief *g_s_*(*t*) about source reliability *θ_s_* is perfectly updated with each new sample of sensory and memory evidence. This is one of the strongest simplifying assumptions of our model, since it is unlikely that observers place equal weight on every sample at every timestep when constructing their belief about *θ_s_*. A simple extension could be introducing stochasticity into the belief-updating process, perhaps by incorporating a softmax function into the update rule defined in Equation 4.3. Another possibility could be that observers differentially attend to or weight evidence based on its *congruence* with the other source, and that this affects the accuracy with which observers construct their belief *g_s_*(*t*). Investigating principled approaches to incorporating asymmetry in belief updating is a promising direction for understanding both the basic mechanisms of reliability-weighted evidence integration, and how they are altered in psychopathology.

## 7 Summary and Conclusion

We have proposed that dynamic reliability-weighted integration of parallel streams of memory and sensory evidence offers a unifying explanation for time-varying effects of expectations on perceptual decisions. To do this, we first discussed how expectations—beliefs about the prior probability of observing a particular outcome—are typically modeled as exerting *static* effects on perceptual decision processes by biasing either the starting point or drift rate of the decision process before any sensory evidence is observed (Section 2). We ended our review by discussing recent findings of *dynamic* effects of expectations on perceptual choice that cannot be explained by static models, and then presented a theoretical framework and computational model suggesting that dynamic effects could be driven by reliability-weighted integration of evidence dynamically retrieved from memory (Sections 3-4). Finally, we showed that our model can reproduce dynamic effects observed across four different experimental tasks, research groups, and neuroimaging modalities, with each effect corresponding to a unique subspace of our broader theoretical framework (Figure 1; Section 5).

These findings suggest that the dynamic reliability-weighted process specified by our model offers a unifying explanation for time-varying effects expectations on perceptual decisions. However, direct empirical evidence supporting our model over multi-stage and/or static alternatives is still lacking, in large part because of how expectation-uncertainty has been omitted from normative models of perceptual decision-making (Section 3). Future work can combine the framework and model we developed in this article with emerging methods for studying decision-making in dynamic and uncertain environments to test the core prediction of our model: the *rate* of evidence accumulation is driven by fluctuations in *relative* reliability of memory and sensory information.

A promising candidate measurement toward this end is the centroparietal positivity (CPP), an event-related EEG signal whose slope has been shown to scale with the strength of both sensory and memory evidence independently (Tsvinev et al., 2025; van Vugt et al., 2019). The conceptual framework we develop in Section 3 predicts that effects of memory sampling should be strongest in fully heterogeneous environments, i.e., those where observers have uncertainty over the true predictive probability of expectation cues, where the cues are manipulated across trials, and where the sensory evidence strength also varies across trials. As suggested above, a behavioral task that dynamically manipulates the *relative* reliability of memory and sensory evidence evidence within a single trial will be necessary to distinguish our parallel integrated model from the serial two-stage models discussed above.

## Acknowledgments

We thank Joachim Vandekerckhove and Mark Steyvers for feedback on an early draft of this manuscript.

## Footnotes

1 Prediction (1) here derives from the *reward-rate* formalization of the speed-accuracy tradeoff, which posits that observers’ objective function is to maximize their total amount of reward (i.e., correct choices) per unit time (e.g., Drugowitsch et al., 2012. In homogeneous environments, reward-rate and SPRT-optimality are equivalent (Moran, 2015); the reward-rate formalization simply adds the ability to make predictions about how task timing ought to impact optimal choice procedures.

2 This dimension of uncertainty roughly corresponds to the description-experience gap as used in the literature on risky, value-based choice (Hertwig and Erev, 2009; Heilbronner and Hayden, 2016).

## References

Afacan-Seref, K., Steinemann, N. A., Blangero, A., & Kelly, S. P. (2018). Dynamic interplay of value and sensory information in high-speed decision making. Curr. Biol., 28 (5), 795–802.e6.

Aitken, F., & Kok, P. (2022). Hippocampal representations switch from errors to predictions during acquisition of predictive associations. Nat. Commun., 13 (1), 3294.

Alister, M., & Evans, N. J. (2026). A diffusion-based framework for modeling systematic, time-varying cognitive processes. Psychol. Rev.

Angelaki, D. E., Gu, Y., & DeAngelis, G. C. (2009). Multisensory integration: Psychophysics, neurophysiology, and computation. Curr. Opin. Neurobiol., 19 (4), 452–458.

Aronowitz, S. (2019). Memory is a modeling system. Mind Lang., 34 (4), 483–502.

Bakkour, A., Palombo, D. J., Zylberberg, A., Kang, Y. H., Reid, A., Verfaellie, M., Shadlen, M. N., & Shohamy, D. (2019). The hippocampus supports deliberation during value-based decisions. Elife, 8.

Banavar, N. V., Noh, S. M., Wahlheim, C. N., Cassidy, B. S., Kirwan, C. B., Stark, C. E., & Bornstein, A. M. (2024). A response time model of the three-choice mnemonic similarity task provides stable, mechanistically interpretable individual-difference measures. Frontiers in human neuroscience, 18, 1379287.

Barendregt, N. W., Gold, J. I., Josić, K., & Kilpatrick, Z. P. (2022). Normative decision rules in changing environments. Elife, 11.

Barker, R. M., Armson, M. J., Diamond, N. B., Liu, Z.-X., Wang, Y., Ryan, J. D., & Levine, B. (2025). Remembrance with gazes passed: Eye movements precede continuous recall of episodic details of real-life events. Cognition, 268 (106380), 106380.

Barnard, G. A. (1946). Sequential tests in industrial statistics. Suppl. J. R. Stat. Soc., 8 (1), 1.

Batchelder, W. H. (2010). Mathematical psychology. Wiley Interdiscip. Rev. Cogn. Sci., 1 (5), 759–765.

Biderman, N., Bakkour, A., & Shohamy, D. (2020). What are memories for? the hippocampus bridges past experience with future decisions. Trends Cogn. Sci., 24 (7), 542–556.

Bogacz, R., Brown, E., Moehlis, J., Holmes, P., & Cohen, J. D. (2006). The physics of optimal decision making: A formal analysis of models of performance in two-alternative forced-choice tasks. Psychol. Rev., 113 (4), 700–765.

Bondy, A. G., Charlton, J. A., Luo, T. Z., Kopec, C. D., Stagnaro, W. M., Venditto, S. J. C., Lynch, L., Janarthanan, S., Oline, S. N., Harris, T. D., & Brody, C. D. (2025). Brain-wide coordination of internal signals during decision-making. bioRxivorg, 2024.08.21.609044.

Bornstein, A. M., Aly, M., Feng, S. F., Turk-Browne, N. B., Norman, K. A., & Cohen, J. D. (2023). Associative memory retrieval modulates upcoming perceptual decisions. Cogn. Affect. Behav. Neurosci.

Bornstein, A. M., & Daw, N. D. (2012). Dissociating hippocampal and striatal contributions to sequential prediction learning. Eur. J. Neurosci., 35 (7), 1011–1023.

Bornstein, A. M., & Daw, N. D. (2013). Cortical and hippocampal correlates of deliberation during model-based decisions for rewards in humans. PLoS Comput. Biol., 9 (12), e1003387.

Britten, K. H., Newsome, W. T., Shadlen, M. N., Celebrini, S., & Movshon, J. A. (1996). A relationship between behavioral choice and the visual responses of neurons in macaque MT. Vis. Neurosci., 13 (1), 87–100.

Brody, C. D., & Hanks, T. D. (2016). Neural underpinnings of the evidence accumulator. Curr. Opin. Neurobiol., 37, 149–157.

Brunton, B. W., Botvinick, M. M., & Brody, C. D. (2013). Rats and humans can optimally accumulate evidence for decision-making. Science, 340 (6128), 95–98.

Callaway, F., Griffiths, T. L., Norman, K. A., & Zhang, Q. (2024). Optimal metacognitive control of memory recall. Psychol. Rev., 131 (3), 781–811.

Carpenter, R. H., & Williams, M. L. (1995). Neural computation of log likelihood in control of saccadic eye movements. Nature, 377 (6544), 59–62.

Chen, J., & Bornstein, A. M. (2024). The causal structure and computational value of narratives. Trends Cogn. Sci., 28 (8), 769–781.

de Lange, F. P., Heilbron, M., & Kok, P. (2018). How do expectations shape perception? Trends Cogn. Sci., 22 (9), 764–779.

de Lange, F. P., Rahnev, D. A., Donner, T. H., & Lau, H. (2013). Prestimulus oscillatory activity over motor cortex reflects perceptual expectations. J. Neurosci., 33 (4), 1400–1410.

Deneve, S. (2012). Making decisions with unknown sensory reliability. Front. Neurosci., 6, 75.

Diaz, J. A., Pisauro, M. A., Delis, I., & Philiastides, M. G. (2024). Prior probability biases perceptual choices by modulating the accumulation rate, rather than the baseline, of decision evidence. Imaging Neurosci. (Camb.), 2, imag–2–00338.

Diederich, A., & Busemeyer, J. R. (2006). Modeling the effects of payoff on response bias in a perceptual discrimination task: Bound-change, drift-rate-change, or two-stage-processing hypothesis. Percept. Psychophys., 68 (2), 194–207.

Diederich, A., & Oswald, P. (2014). Sequential sampling model for multiattribute choice alternatives with random attention time and processing order. Front. Hum. Neurosci., 8, 697.

Diederich, A., & Oswald, P. (2016). Multi-stage sequential sampling models with finite or infinite time horizon and variable boundaries. J. Math. Psychol., 74, 128–145.

Drugowitsch, J., Moreno-Bote, R., Churchland, A. K., Shadlen, M. N., & Pouget, A. (2012). The cost of accumulating evidence in perceptual decision making. J. Neurosci., 32 (11), 3612–3628.

Dunovan, K. E., Tremel, J. J., & Wheeler, M. E. (2014). Prior probability and feature predictability interactively bias perceptual decisions. Neuropsychologia, 61, 210–221.

Dunovan, K. E., & Wheeler, M. E. (2018). Computational and neural signatures of pre and post-sensory expectation bias in inferior temporal cortex. Sci. Rep., 8 (1), 13256.

Edwards, W. (1965). Optimal strategies for seeking information: Models for statistics, choice reaction times, and human information processing. J. Math. Psychol., 2 (2), 312–329.

Fang, M., Mao, J., Donner, T. H., & Stocker, A. A. (2026). The resource-rational dynamics of evidence accumulation. bioRxiv, 2026.04.15.718716.

Fetsch, C. R., Kiani, R., Newsome, W. T., & Shadlen, M. N. (2014). Effects of cortical microstimulation on confidence in a perceptual decision. Neuron, 84 (1), 239.

Fetsch, C. R., Pouget, A., DeAngelis, G. C., & Angelaki, D. E. (2011). Neural correlates of reliability-based cue weighting during multisensory integration. Nat. Neurosci., 15 (1), 146–154.

Findling, C., Hubert, F., International Brain Laboratory, Acerbi, L., Benson, B., Benson, J., Birman, D., Bonacchi, N., Buchanan, E. K., Bruijns, S., Carandini, M., Catarino, J. A., Chapuis, G. A., Churchland, A. K., Dan, Y., Davatolhagh, F., DeWitt, E. E. J., Engel, T. A., Fabbri, M.,... Pouget, A. (2025). Brain-wide representations of prior information in mouse decision-making. Nature, 645 (8079), 192–200.

Finn, E. S., Corlett, P. R., Chen, G., Bandettini, P. A., & Constable, R. T. (2018). Trait paranoia shapes inter-subject synchrony in brain activity during an ambiguous social narrative. Nat. Commun., 9 (1), 2043.

Forstmann, B. U., Ratcliff, R., & Wagenmakers, E.-J. (2016). Sequential sampling models in cognitive neuroscience: Advantages, applications, and extensions. Annu. Rev. Psychol., 67, 641–666.

Frank, D., Moratti, S., Hellerstedt, R., Sarnthein, J., Li, N., Horn, A., Imbach, L., Stieglitz, L., Gil-Nagel, A., Toledano, R., Friston, K. J., & Strange, B. A. (2026). Human hippocampal ripples tune cortical responses based on predicted uncertainty. Nat. Neurosci., 29, 1987–1998.

Frazier, P., & Yu, A. J. (2007). Sequential hypothesis testing under stochastic deadlines. Adv. Neural Inf. Process. Syst., 465–472.

Gläscher, J., Daw, N., Dayan, P., & O’Doherty, J. P. (2010). States versus rewards: Dissociable neural prediction error signals underlying model-based and model-free reinforcement learning. Neuron, 66 (4), 585–595.

Glaze, C. M., Kable, J. W., & Gold, J. I. (2015). Normative evidence accumulation in unpredictable environments. Elife, 4.

Gold, J. I., & Shadlen, M. N. (2002). Banburismus and the brain: Decoding the relationship between sensory stimuli, decisions, and reward. Neuron, 36 (2), 299–308.

Gold, J. I., & Shadlen, M. N. (2007). The neural basis of decision making. Annu. Rev. Neurosci., 30, 535–574.

Hachisuka, A., Shor, J. D., Liu, X. C., Friedman, D., Dugan, P., Saez, I., Panov, F. E., Wang, Y., Doyle, W., Devinsky, O., Oermann, E. K., & He, B. J. (2026). Neural and computational mechanisms underlying one-shot perceptual learning in humans. Nat. Commun., 17 (1), 1204.

Hanks, T. D., Mazurek, M. E., Kiani, R., Hopp, E., & Shadlen, M. N. (2011). Elapsed decision time affects the weighting of prior probability in a perceptual decision task. J. Neurosci., 31 (17), 6339–6352.

Hanks, T. D., & Summerfield, C. (2017). Perceptual decision making in rodents, monkeys, and humans. Neuron, 93 (1), 15–31.

Hannula, D. E., Althoff, R. R., Warren, D. E., Riggs, L., Cohen, N. J., & Ryan, J. D. (2010). Worth a glance: Using eye movements to investigate the cognitive neuroscience of memory. Front. Hum. Neurosci., 4, 166.

Heilbronner, S. R., & Hayden, B. Y. (2016). The description-experience gap in risky choice in nonhuman primates. Psychon. Bull. Rev., 23 (2), 593–600.

Hertwig, R., & Erev, I. (2009). The description-experience gap in risky choice. Trends Cogn. Sci., 13 (12), 517–523.

Herweg, N. A., Solomon, E. A., & Kahana, M. J. (2020). Theta oscillations in human memory. Trends Cogn. Sci., 24 (3), 208–227.

Huang, Y., Friesen, A. L., Hanks, T. D., Shadlen, M. N., & Rao, R. P. N. (2012). How prior probability influences decision making: A unifying probabilistic model. Adv. Neural Inf. Process. Syst., 25 (1).

Jacobs, J. (2014). Hippocampal theta oscillations are slower in humans than in rodents: Implications for models of spatial navigation and memory. Philos. Trans. R. Soc. Lond. B Biol. Sci., 369 (1635), 20130304.

Kaelbling, L. P., Littman, M. L., & Cassandra, A. R. (1998). Planning and acting in partially observable stochastic domains. Artif. Intell., 101 (1-2), 99–134.

Kaper, R., & Peters, M. (2026). Stimulus familiarity shapes hierarchical structure learning and metacognitive dynamics. PsyArXiv.

Kelly, S. P., Corbett, E. A., & O’Connell, R. G. (2021). Neurocomputational mechanisms of prior-informed perceptual decision-making in humans. Nat Hum Behav, 5 (4), 467–481.

Kerrén, C., Linde-Domingo, J., Hanslmayr, S., & Wimber, M. (2018). An optimal oscillatory phase for pattern reactivation during memory retrieval. Curr. Biol., 28 (21), 3383–3392.e6.

Kerrén, C., Michelmann, S., & Doeller, C. F. (2026). Hippocampal ripples initiate cortical dimensionality expansion for memory retrieval. Nat. Commun., 17, 6677.

Khodadadi, A., Fakhari, P., & Busemeyer, J. R. (2014). Learning to maximize reward rate: A model based on semi-markov decision processes. Front. Neurosci., 8, 101.

Khoudary, A., Bornstein, A., & Peters, M. (2025a). Investigating implicit and explicit expectations in perceptual decision making. 47.

Khoudary, A., Peters, M. A. K., & Bornstein, A. M. (2025b). Reasoning goals and representational decisions in computational cognitive neuroscience: Lessons from the drift diffusion model. Eur. J. Neurosci., 61 (7), e70098.

Kiani, R., Churchland, A. K., & Shadlen, M. N. (2013). Integration of direction cues is invariant to the temporal gap between them. J. Neurosci., 33 (42), 16483–16489.

Kiani, R., Corthell, L., & Shadlen, M. N. (2014). Choice certainty is informed by both evidence and decision time. Neuron, 84 (6), 1329–1342.

Kilpatrick, Z. P., Holmes, W. R., Eissa, T. L., & Josić, K. (2019). Optimal models of decision-making in dynamic environments. Curr. Opin. Neurobiol., 58, 54–60.

Kok, P., Brouwer, G. J., van Gerven, M. A. J., & de Lange, F. P. (2013). Prior expectations bias sensory representations in visual cortex. J. Neurosci., 33 (41), 16275–16284.

Kok, P., Jehee, J. F. M., & de Lange, F. P. (2012). Less is more: Expectation sharpens representations in the primary visual cortex. Neuron, 75 (2), 265–270.

Kragel, J. E., VanHaerents, S., Templer, J. W., Schuele, S., Rosenow, J. M., Nilakantan, A. S., & Bridge, D. J. (2020). Hippocampal theta coordinates memory processing during visual exploration. Elife, 9 (e52108).

Krajbich, I., Armel, C., & Rangel, A. (2010). Visual fixations and the computation and comparison of value in simple choice. Nat. Neurosci., 13 (10), 1292–1298.

Kumle, L., Nobre, A. C., & Draschkow, D. (2025). Sensorimnemonic decisions: Choosing memories versus sensory information. Trends Cogn. Sci.

Landy, M. S., Banks, M. S., & Knill, D. C. (2011). Ideal-observer models of cue integration. In J. Trommershauser, K. Kording, & M. S. Landy (Eds.), Sensory cue integration. Oxford University Press.

Link, S. W. (1975). The relative judgment theory of two choice response time. J. Math. Psychol., 12 (1), 114–135.

Liu, A., Schartner, M., International Brain Laboratory, & Fiete, I. (2025). How learned expectations shape brain-wide responses. bioRxiv, 2025.12.15.694430.

Malhotra, G., Leslie, D. S., Ludwig, C. J. H., & Bogacz, R. (2017). Overcoming indecision by changing the decision boundary. J. Exp. Psychol. Gen., 146 (6), 776–805.

Malhotra, G., Leslie, D. S., Ludwig, C. J. H., & Bogacz, R. (2018). Time-varying decision boundaries: Insights from optimality analysis. Psychon. Bull. Rev., 25 (3), 971–996.

Mazor, M., Moran, R., & Press, C. (2026). Beliefs about perception shape perceptual inference: An ideal observer model of detection. Psychol. Rev., 133 (2), 271–295.

Moran, R. (2015). Optimal decision making in heterogeneous and biased environments. Psychon. Bull. Rev., 22 (1), 38–53.

Mulder, M. J., Wagenmakers, E.-J., Ratcliff, R., Boekel, W., & Forstmann, B. U. (2012). Bias in the brain: A diffusion model analysis of prior probability and potential payoff. J. Neurosci., 32 (7), 2335–2343.

Newsome, W. T., Britten, K. H., & Movshon, J. A. (1989). Neuronal correlates of a perceptual decision. Nature, 341 (6237), 52–54.

Nicholas, J., Chen, S., & Mattar, M. G. (2026). Flexible decisions arise from resource-rational memory sampling. bioRxiv, 2026.06. 15.732446.

Nicholas, J., & Mattar, M. G. (2026). Episodic memory facilitates flexible decision-making via access to detailed events. *Nat*. Hum. Behav., 1–17.

Noh, S. M., Singla, U. K., Bennett, I. J., & Bornstein, A. M. (2023). Memory precision and age differentially predict the use of decision-making strategies across the lifespan. Sci. Rep., 13 (1), 17014.

O’Connell, R. G., Dockree, P. M., & Kelly, S. P. (2012). A supramodal accumulation-to-bound signal that determines perceptual decisions in humans. Nat. Neurosci., 15 (12), 1729–1735.

O’Connell, R. G., Shadlen, M. N., Wong-Lin, K., & Kelly, S. P. (2018). Bridging neural and computational viewpoints on perceptual decision-making. Trends Neurosci., 41 (11), 838–852.

Oliva, A., & Torralba, A. (2007). The role of context in object recognition. Trends Cogn. Sci., 11 (12), 520– 527.

Palmer, J., Huk, A. C., & Shadlen, M. N. (2005a). The effect of stimulus strength on the speed and accuracy of a perceptual decision. J. Vis., 5 (5), 376–404.

Palmer, J., McKinley, M. K., Mazurek, M., & Shadlen, M. N. (2005b). Effect of prior probability on choice and response time in a motion discrimination task. J. Vis., 5 (8), 235–235.

Polanía, R., Krajbich, I., Grueschow, M., & Ruff, C. C. (2014). Neural oscillations and synchronization differentially support evidence accumulation in perceptual and value-based decision making. Neuron, 82 (3), 709–720.

Portides, D. (2021). Idealization and abstraction in scientific modeling. Synthese, 198 (24), 5873–5895.

Poskanzer, C., & Aly, M. (2023). Switching between external and internal attention in hippocampal networks. J. Neurosci., 43 (38), 6538–6552.

Press, C., Kok, P., & Yon, D. (2020). The perceptual prediction paradox. Trends Cogn. Sci., 24 (1), 13–24.

Rao, R. P. N. (2010). Decision making under uncertainty: A neural model based on partially observable markov decision processes. Front. Comput. Neurosci., 4, 146.

Rao, V., DeAngelis, G. C., & Snyder, L. H. (2012). Neural correlates of prior expectations of motion in the lateral intraparietal and middle temporal areas. J. Neurosci., 32 (29), 10063–10074.

Ratcliff, R. (1978). A theory of memory retrieval. Psychol. Rev., 85 (2), 59–108.

Ratcliff, R. (1980). A note on modeling accumulation of information when the rate of accumulation changes over time. J. Math. Psychol., 21 (2), 178–184.

Ratcliff, R., & McKoon, G. (2008). The diffusion decision model: Theory and data for two-choice decision tasks. Neural Comput., 20 (4), 873–922.

Ratcliff, R., & Rouder, J. N. (1998). Modeling response times for two-choice decisions. Psychol. Sci., 9 (5), 347–356.

Ratcliff, R., Smith, P. L., Brown, S. D., & McKoon, G. (2016). Diffusion decision model: Current issues and history. Trends Cogn. Sci., 20 (4), 260–281.

Ratcliff, R., & Tuerlinckx, F. (2002). Estimating parameters of the diffusion model: Approaches to dealing with contaminant reaction times and parameter variability. Psychon. Bull. Rev., 9 (3), 438–481.

Rittershofer, K., Wang, Y., Eimer, M., Kok, P., Yon, D., & Clare Press. (2026). Paradoxical influences of prediction are resolved across time. *bioRxiv*, 2026.03.05.709935.

Roweis, S., & Ghahramani, Z. (1998). A unifying review of linear gaussian models. Neural Comput., 11 (2), 34.

Rungratsameetaweemana, N., Itthipuripat, S., Salazar, A., & Serences, J. T. (2018). Expectations do not alter early sensory processing during perceptual decision-making. J. Neurosci., 38 (24), 5632–5648.

Rungratsameetaweemana, N., & Serences, J. T. (2019). Dissociating the impact of attention and expectation on early sensory processing. Curr. Opin. Psychol., 29, 181–186.

Seger, S. E., Kriegel, J. L. S., Lega, B. C., & Ekstrom, A. D. (2023). Memory-related processing is the primary driver of human hippocampal theta oscillations. Neuron, 111 (19), 3119–3130.e4.

Senoussi, M., Galas, L., Busch, N. A., & Dugué, L. (2025). Theta-rhythmic attentional exploration of space. bioRxiv, 2025.08.16.670674.

Seriès, P., & Seitz, A. R. (2013). Learning what to expect (in visual perception). Front. Hum. Neurosci., 7, 668.

Shadlen, M. N., & Shohamy, D. (2016). Decision making and sequential sampling from memory. Neuron, 90 (5), 927–939.

Shi, Y.-L., Zeraati, R., International Brain Laboratory, Levina, A., & Engel, T. A. (2025). Brain-wide organization of intrinsic timescales at single-neuron resolution. bioRxivorg.

Shushruth, S., Zylberberg, A., & Shadlen, M. N. (2022). Sequential sampling from memory underlies action selection during abstract decision-making. Curr. Biol., 32 (9), 1949–1960.e5.

Simen, P., Contreras, D., Buck, C., Hu, P., Holmes, P., & Cohen, J. D. (2009). Reward rate optimization in two-alternative decision making: Empirical tests of theoretical predictions. J. Exp. Psychol. Hum. Percept. Perform., 35 (6), 1865–1897.

Srivastava, V., Feng, S. F., Cohen, J. D., Leonard, N. E., & Shenhav, A. (2017). A martingale analysis of first passage times of time-dependent wiener diffusion models. J. Math. Psychol., 77, 94–110.

Stone, M. (1960). Models for choice-reaction time. Psychometrika, 25 (3), 251–260.

Summerfield, C., & de Lange, F. P. (2014). Expectation in perceptual decision making: Neural and computational mechanisms. Nat. Rev. Neurosci., 15 (11), 745–756.

Tarder-Stoll, H., Sekeres, M. J., Levine, B., & Moscovitch, M. (2026). Adaptive episodic memory: How multiple memory representations drive behavior in humans and nonhumans. Physiol. Rev., 106 (2), 841–889.

Tavares, G., Perona, P., & Rangel, A. (2017). The attentional drift diffusion model of simple perceptual decision-making. Front. Neurosci., 11, 468.

Ter Wal, M., Linde-Domingo, J., Lifanov, J., Roux, F., Kolibius, L. D., Gollwitzer, S., Lang, J., Hamer, H., Rollings, D., Sawlani, V., Chelvarajah, R., Staresina, B., Hanslmayr, S., & Wimber, M. (2021). Theta rhythmicity governs human behavior and hippocampal signals during memory-dependent tasks. Nat. Commun., 12 (1), 7048.

Tsvinev, A., Pilipenko, A., & Samaha, J. (2025). A common neural signal of evidence accumulation for perceptual and mnemonic decisions. bioRxiv, 2025.11. 13.688140.

van Ravenzwaaij, D., Mulder, M. J., Tuerlinckx, F., & Wagenmakers, E.-J. (2012). Do the dynamics of prior information depend on task context? an analysis of optimal performance and an empirical test. Front. Psychol., 3, 132.

van Vugt, M. K., Beulen, M. A., & Taatgen, N. A. (2019). Relation between centro-parietal positivity and diffusion model parameters in both perceptual and memory-based decision making. Brain Res., 1715, 1–12.

Verschooren, S., Dahl, M. J., Aly, M., & Mittner, M. (2026). Transition dynamics of external and internal attention across on-task and off-task states. Nat. Rev. Psychol., 1–17.

Vickers, D. (1970). Evidence for an accumulator model of psychophysical discrimination. Ergonomics, 13 (1), 37–58.

Vivekananda, U., Bush, D., Bisby, J. A., Baxendale, S., Rodionov, R., Diehl, B., Chowdhury, F. A., McEvoy, A. W., Miserocchi, A., Walker, M. C., & Burgess, N. (2021). Theta power and theta-gamma coupling support long-term spatial memory retrieval. Hippocampus, 31 (2), 213–220.

Wald, A., & Wolfowitz, J. (1948). Optimum character of the sequential probability ratio test. Ann. Math. Stat., 19 (3), 326–339.

Wald, A. (1945). Sequential tests of statistical hypotheses. Ann. Math. Stat., 16 (2), 117–186.

Wang, S., Feng, S. F., & Bornstein, A. M. (2021). Mixing memory and desire: How memory reactivation supports deliberative decision-making. Wiley Interdiscip. Rev. Cogn. Sci., 13 (2), e1581.

Wert, S., Seidle, A., Rissman, J., & Knowlton, B. J. (2025). The temporal evolution of implicit bias in perceptual decision-making. CogSci, 47 (0).

Yang, X., & Krajbich, I. (2023). A dynamic computational model of gaze and choice in multi-attribute decisions. Psychol. Rev., 130 (1), 52–70.

Yoo, J., & Bornstein, A. (2024). Temporal dynamics of model-based control reveal arbitration between multiple task representations. *PsyArXiv*.

Zeithamova, D., Schlichting, M. L., & Preston, A. R. (2012). The hippocampus and inferential reasoning: Building memories to navigate future decisions. Front. Hum. Neurosci., 6, 70.

Zhang, H., Watrous, A. J., Patel, A., & Jacobs, J. (2018). Theta and alpha oscillations are traveling waves in the human neocortex. Neuron, 98 (6), 1269–1281.e4.

Zhou, Z., & Geng, J. J. (2024). Learned associations serve as target proxies during difficult but not easy visual search. Cognition, 242 (105648), 105648.

Zubair, H. N., Stangl, M., Topalovic, U., Inman, C., Seeber, M., Hiller, S., Rao, V. R., Halpern, C. H., Eliashiv, D., Fried, I., & Suthana, N. (2026). Eye movements reflect memory-related theta activity in the human brain. PLoS Biol., 24 (3), e3003695.

Zylberberg, A., Bakkour, A., Shohamy, D., & Shadlen, M. N. (2024). Value construction through sequential sampling explains serial dependencies in decision making. Elife, 13 (RP96997), RP96997.

Zylberberg, A., Fetsch, C. R., & Shadlen, M. N. (2016). The influence of evidence volatility on choice, reaction time and confidence in a perceptual decision. Elife, 5.

