## Supplementary Materials for "Memory retrieval explains dynamic effects of expectations on perceptual decisions"

#### S1 Qualitative comparison of normative starting point models

Visualizing prescriptions made by the starting point models of Edwards (1965) and Link (1975) demonstrates the exponential scaling of optimal starting points by the signal-to-noise ratio of sensory evidence in the environment.

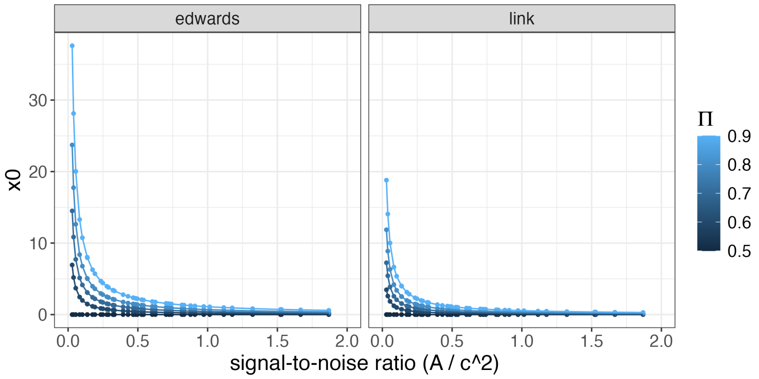


Figure S1: Optimal starting points on the models of Edwards (1965) and Link (1975).

#### S2 Sequential effects analysis

This section contains details about the two studies used to demonstrate sequential effects of expectations on RTs. For each study, we provide a brief summary, followed by the regression model specification and a full summary table of fitted coefficients. Reaction times were log-transformed and z-scored within subject prior to being entered into the regression. All predictors were defined categorically to maximize the descriptive ability of each model (i.e., we did not force reaction times to scale numerically with cue predictiveness), except for sensory evidence strength and time on task, which were treated as continuous. Models were fit using the `lm()` function in `R 4.4.2`, and summary tables were produced using the `tbl_regression()` function from the `gtsummary` package.

##### S2.1 Diaz, Pisauro et al. (2024): instructed expectations

The data from Diaz, Pisauro et al. (2024)were generated by human observers (n=16) performing a face/car discrimination task in a fully heterogeneous environment with explicitly instructed $\Pi_{cue}$. Each trial began with a brief (750ms) presentation of one of three possible cues—30F, 50F, or 70F—overtly instructing observers on the trial-level probability that the correct answer will be face. A noise-degraded stimulus was then briefly (50ms) presented, and observers had up to 1250ms to indicate whether the stimulus contained an image of a car or a face. Stimuli were presented at moderate levels of signal strength (coherence = 32.5% and 37.5%), and observers received no feedback on the accuracy of their choice. Each cue was presented 360 times for a total of 1080 trials per subject.

We modeled trial-by-trial transformed reaction times ($RT_t$) using the following formula:

$$\mathrm{RT}_t \sim \mathrm{Accuracy}_t + \mathrm{Validity}_t + \mathrm{PreviousCue}_t + \mathrm{PreviousTarget}_t + \mathrm{PreviousResponse}_t \tag{S1}$$

The results of this model are displayed in Table S2.1 below. All factors canonically known to modify RTs—accuracy, cue validity, and the accuracy of the previous response—exhibited significant main effects ($\beta_{accuracy} = -0.23$, $\beta_{valid} = -0.12$, $\beta_{invalid} = 0.05$, $\beta_{prev\_correct} = -0.03$). Crucially, switching cues across trials slowed RTs ($\beta_{prev\_cue} = 0.03$) but switching *targets* across trials had no effect on RTs ($\beta_{prev\_target} = 0.00$). The null effect of sequential targets is visualized in Figure **??**.

| **Predictor** | **Beta** | **SE** | **t-statistic** | **95% CI** | **p-value** |
|---|---|---|---|---|---|
| accuracy | | | | | |
| correct | -0.20 | 0.011 | -17.4 | -0.22, -0.17 | <0.001 |
| incorrect | — | — | — | — | |
| validity | | | | | |
| valid | -0.12 | 0.010 | -12.0 | -0.14, -0.10 | <0.001 |
| invalid | 0.05 | 0.010 | 5.23 | 0.03, 0.07 | <0.001 |
| neutral | — | — | — | — | |
| prev_cue | | | | | |
| different | 0.03 | 0.008 | 3.86 | 0.01, 0.04 | <0.001 |
| same | — | — | — | — | |
| prev_target | | | | | |
| different | 0.00 | 0.007 | 0.481 | -0.01, 0.02 | 0.6 |
| same | — | — | — | — | |
| prev_response | | | | | |
| correct | 0.00 | 0.011 | 0.290 | -0.02, 0.03 | 0.8 |
| error | — | — | — | — | |
| logTrial | -0.23 | 0.007 | -31.2 | -0.25, -0.22 | <0.001 |

No. Obs. = 16,696; Sigma = 0.915; Statistic = 221; p-value = <0.001; df = 7; Residual df = 16,688

#### S2.2 Bornstein et al. (2023): learned expectations

Data from Bornstein et al. (2023) were generated by human observers (n=33) performing a category-level discrimination task of noise-degraded, perceptually-similar grayscale images (i.e., discriminating Face A from Face B). $\Pi_{cue}$ was held constant for within-category stimuli, such that one category was always more predictable than the other, but each stimulus within the category had a unique cue (colored fractal). The predictability of each category (faces and scenes) was manipulated across blocks, such that each observer learned distinct cues on the range $\Pi_{cue} = \{0.5, 0.6, 0.7, 0.8\}$. Within each block, observers had 30 trials per cue (120 trials total) to learn the predictive relationship of each unique colored fractal and its corresponding grayscale image; this learning procedure ensures that observers have a source-related uncertainty in their expectations. Sensory evidence was presented at "low" and "high" signal strength, corresponding to 65% and 85% discrimination accuracy in the absence of cues. Coherence was yoked to stimulus category, such that stimuli with more predictive cues were always presented with low coherence whereas stimuli with less predictive cues were always presented with high coherence. Each trial began with a brief (750ms) presentation of one of the four fractal cues, followed by a variable delay of 4000, 6000, or 8000ms. Observers were then shown a 60Hz "stream" of within-category images, and they had up to 3000ms to report which image was presented more frequently. Immediately after responding, the correct image appeared on screen surrounded by a colored box indicating whether the response was correct (green) or incorrect (red). There was no time penalty for incorrect responses. Each block contained 20 decision trials per cue, such that each observer contributed 160 trials total.

We modeled trial-by-trial transformed reaction times ($RT_t$) using the following formula:

$$\mathrm{RT}_t \sim \mathrm{cueLevel}_t * \mathrm{validity}_t + \mathrm{Accuracy}_t + \mathrm{PreviousCue}_t + \mathrm{PreviousTarget}_t + \mathrm{PreviousResponse}_t \quad \text{(S2)}$$

This model is slightly more complex than the others due to the nature of the task. Bornstein et al. (2023) measured effects of four different cue levels for each subject, whereas the other studies only had one level of cue. Accordingly, it was necessary for cue validity to interact with the strength of the cue for this dataset, whereas the other datasets could use cue validity alone as the predictor. We treated `cueLevel` as a categorical variable to ensure that any variability due to errors in learning would be captured by the model (as opposed to enforcing a strictly linear effect of the cue's true predictive probability).

The results of this model are displayed in Table S2.1 below. Choice accuracy, cue level, and previous cue all exhibited significant main effects on reaction times ($\beta_{accuracy} = 0.16$, $\beta_{cueLevel\_0.5} = 0.21$, $\beta_{cueLevel\_0.7} = -0.06$, $\beta_{prev\_cue} = 0.06$). The positive relationship between correct responses and reaction times is due to including both "early" and "late" responses in the same regression; the canonical speed-accuracy relationship is observed when $\beta$ values are estimated independently for "early" and "late" responses (Bornstein et al., 2023).

| Predictor | Beta | SE | t-statistic | 95% CI | p-value |
|---|---|---|---|---|---|
| Accuracy | | | | | |
| correct | 0.16 | 0.016 | 9.95 | 0.13, 0.19 | <0.001 |
| incorrect | — | — | — | — | |
| cueLevel | | | | | |
| 0.5 | 0.21 | 0.025 | 8.16 | 0.16, 0.26 | <0.001 |
| 0.6 | 0.05 | 0.026 | 1.79 | 0.00, 0.10 | 0.074 |
| 0.7 | -0.05 | 0.027 | -2.03 | -0.11, 0.00 | 0.043 |
| 0.8 | — | — | — | — | |
| validity | | | | | |
| invalid | 0.02 | 0.016 | 1.36 | -0.01, 0.05 | 0.2 |
| valid | — | — | — | — | |
| prev_cue | | | | | |
| different | 0.06 | 0.017 | 3.15 | 0.02, 0.09 | 0.002 |
| same | — | — | — | — | |
| prev_target | | | | | |
| different | -0.01 | 0.020 | -0.343 | -0.05, 0.03 | 0.7 |
| same | — | — | — | — | |
| prev_response | | | | | |
| correct | -0.01 | 0.015 | -0.926 | -0.04, 0.02 | 0.4 |
| error | — | — | — | — | |
| log(Trial) | -0.24 | 0.085 | -2.81 | -0.41, -0.07 | 0.005 |
| cueLevel * validity | | | | | |
| 0.5 * invalid | -0.03 | 0.026 | -1.24 | -0.08, 0.02 | 0.2 |
| 0.6 * invalid | 0.00 | 0.026 | -0.051 | -0.05, 0.05 | >0.9 |
| 0.7 * invalid | 0.02 | 0.027 | 0.571 | -0.04, 0.07 | 0.6 |

No. Obs. = 4,686; Sigma = 0.973; Statistic = 18.9; p-value = <0.001; df = 12; Residual df = 4,673

#### S2.3 Rungratsameetaweemana, Itthipuripat et al. (2018)effect

This section shows that the main-text results reported in Figure 5 are preserved when averaging across all levels of evidence encoding noise $\epsilon$ (Figure S2A) and when splitting up the time-varying weights by low and high flicker rate (Figure S2B).

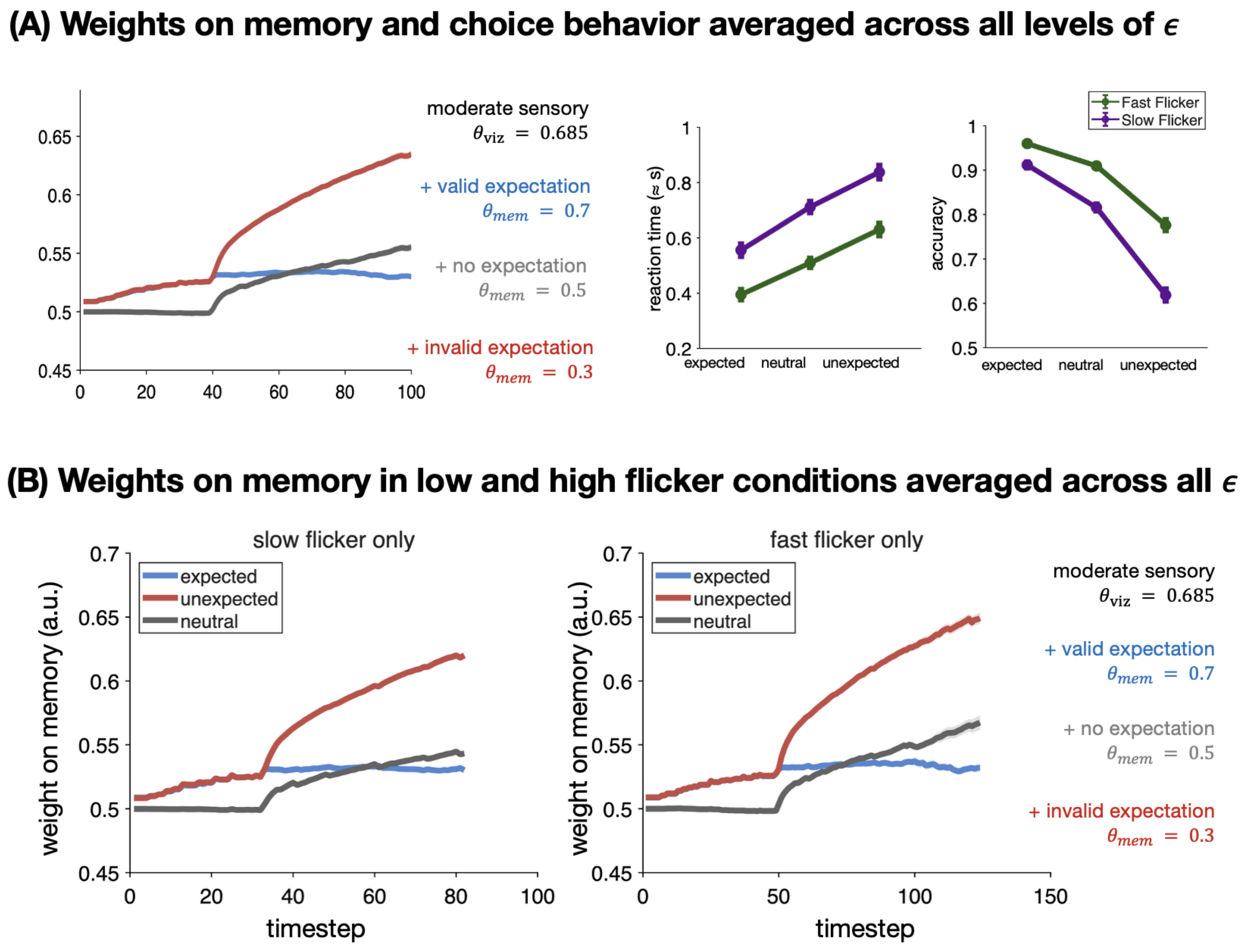


Figure S2: (A) The qualitative effects produced by our model are preserved when averaging over evidence encoding noise $\epsilon$. (B) Dynamic weights on memory generated in the low (33Hz) and high (50Hz) flicker rate simulations of Rungratsameetaweemana, Itthipuripat et al. (2018). Simulations with low flicker rates used $\gamma \in \{4, 6, 8, 10, 12, 20\}$ corresponding to activity at 8.25, 5.5, 4.125, 3.3, 2.5, and 1.5Hz, and simulations with high flicker rates used $\gamma \in \{4, 6, 8, 10, 12, 20\}$ to generate memory samples at 8.3, 6.25, 4.16, 3.5, 2.5, and 1.5Hz. Threshold values $z$ were fixed across conditions within a participant, but varied across participants; data were generated using $z \in \{3, 4, 5\}$. Weights in each condition cease being estimated once the decision variable hits threshold, leading to an increased weighting of invalid expectations at the group level.

#### S3 Effect of cue-stimulus interval duration on Bornstein et al. (2023) effect

Figure S3 shows that the uncertainty-adaptive anticipatory sampling effect displayed in Figure 7 is preserved for both the medium (top) and long (bottom) cue-stimulus interval durations. Visual evidence onset is indicated by the solid vertical line; note difference in x-axes across plots.

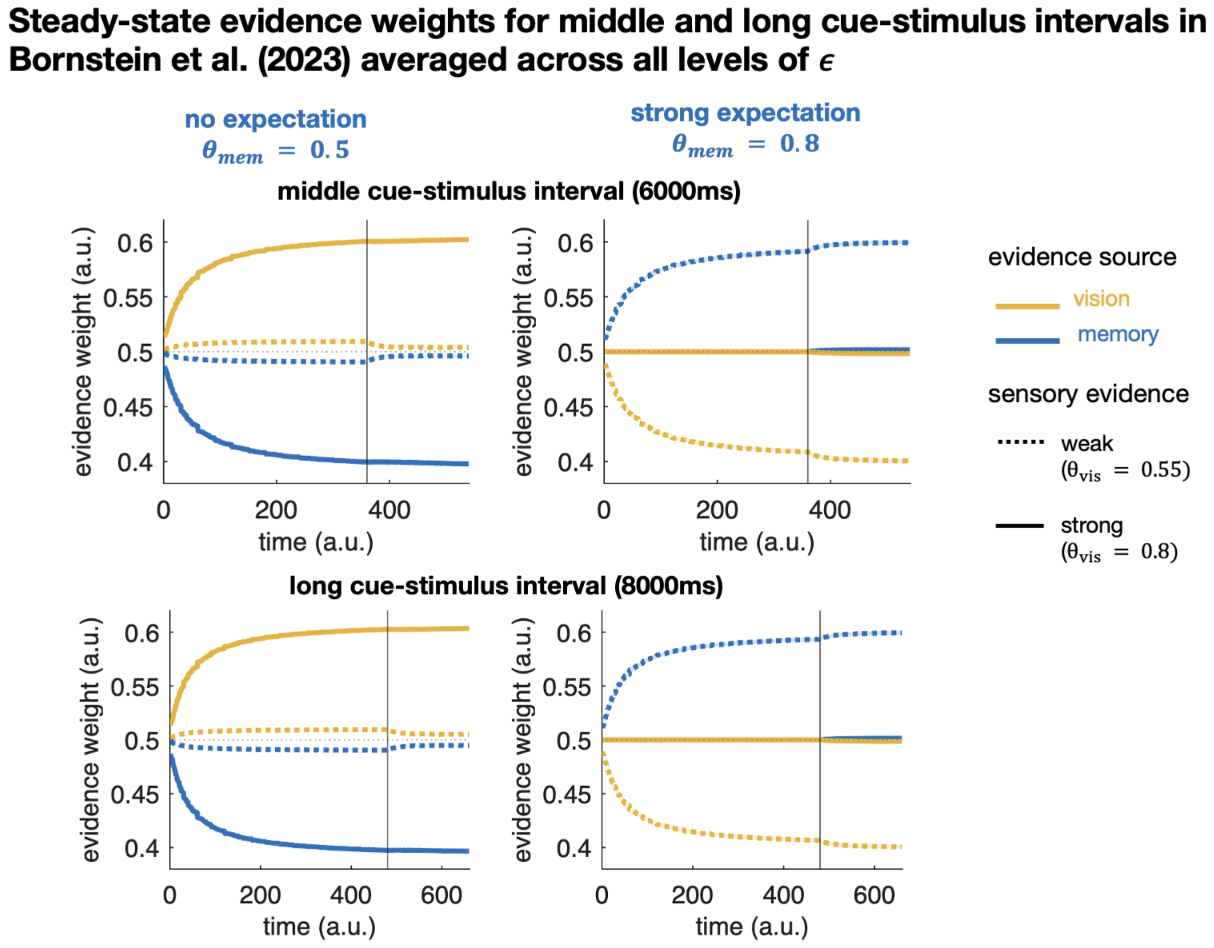


Figure S3: Steady-state evidence weights displaying the uncertainty-adaptive sampling effect observed by Bornstein et al. (2023) at intermediate (top) and long (bottom) cue-stimulus interval durations. Standard error across n=50 steady-state timeseries is plotted but not visible due to its magnitude.

### S4 Hachisuka, Shor, Liu et al. (2026) effect

This section shows that the main-text results in Figure 8 effect are preserved when averaging across all levels of evidence encoding noise $\epsilon$.

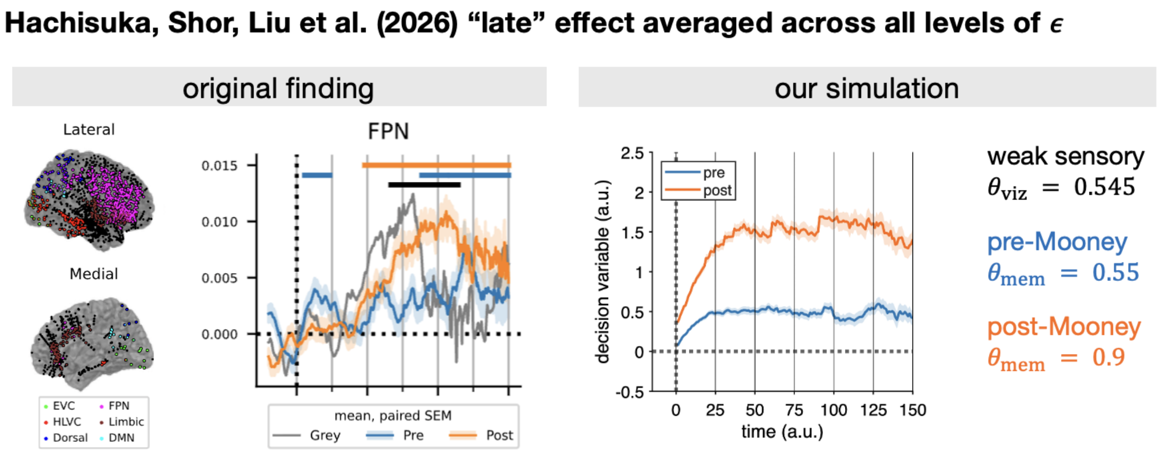


Figure S4: Timeseries from run-averaged simulations of Hachisuka, Shor, Liu et al. (2026). Solid lines are cue-level decision variables averaged across the mean decision variable for each combination of $\gamma$, $z$, and $\epsilon$; shaded areas reflect standard error of the mean (n=100).
